# Improving personalized prediction of cancer prognoses with clonal evolution models

**DOI:** 10.1101/761510

**Authors:** Yifeng Tao, Ashok Rajaraman, Xiaoyue Cui, Ziyi Cui, Jesse Eaton, Hannah Kim, Jian Ma, Russell Schwartz

**Affiliations:** Computational Biology Department, School of Computer Science, Carnegie Mellon University, Pittsburgh, PA 15213, USA; Department of Biological Sciences, Carnegie Mellon University, Pittsburgh, PA 15213, USA

**Author notes:** Correspondence (R.S.); (J.M.).

**Keywords:** Cancer phylogenetics, Mutator phenotypes, Evolutionary features, Cox regression, Prognostic prediction

## Abstract

Cancer occurs via an accumulation of somatic genomic alterations in a process of clonal evolution. There has been intensive study of potential causal mutations driving cancer development and progression. However, much recent evidence suggests that tumor evolution is normally driven by a variety of mechanisms of somatic hypermutability, known as mutator phenotypes, which act in different combinations or degrees in different cancers. Here we explore the question of how and to which degree different mutator phenotypes act in a cancer predict its future progression. We develop a computational paradigm using evolutionary tree inference (tumor phylogeny) algorithms to derive features quantifying single-tumor mutational preferences, followed by a machine learning framework to identify key features predictive of progression. We build phylogenies tracing the evolution of subclones of cells in tumor tissues using a variety of somatic genomic alterations, including single nucleotide variations, copy number alterations, and structural variations. We demonstrate that mutation preference features derived from the phylogenies are predictive of clinical outcomes of cancer progression – overall survival and disease-free survival – based on the analyses on breast invasive carcinoma, lung adenocarcinoma, and lung squamous cell carcinoma. We further show that mutational phenotypes have predictive power even after accounting for traditional clinical and driver-centric predictors of progression. These results confirm the power of mutational phenotypes as an independent class of predictive biomarkers and suggest a strategy for enhancing the predictive power of conventional clinical or driver-centric genomic features.

## 1 Background

Cancers are typically caused by somatic genomic alterations accumulating under the forces of evolutionary diversification and selection that ultimately lead to uncontrolled cell growth [1]. In most cases, cancer progression is accelerated by somatic hypermutability, where defects in DNA replication and/or repair mechanisms cause the rapid acquisition of mutations across generations of cell growth [2]. Tumor cell populations thus typically undergo substantial genetic diversification over time, most of it likely selectively neutral but some with phenotypic effects [3], resulting in profound intra-tumor heterogeneity (ITH) [4], i.e., cell-to-cell variation in terms of genetic makeup. Such heterogeneity in turn creates an opportunity for selection of mutations that promote uncontrolled cell growth, leading ultimately to tumor growth and potentially subsequent metastasis and patient mortality [1, 5]. This process of evolutionary diversification and selection further underlies the development of cancer resistance to therapeutics [6]. Understanding the processes of somatic evolution that act in cancers is thus crucial to understanding why some precancerous lesions progress to cancer while others do not, why some cancers are highly aggressive while others are indolent, and why some respond robustly to treatment and others do not [7].

One of the key insights into cancer progression to derive from high-throughput sequencing studies is that mechanisms of somatic evolution can differ widely across cancer types. Mechanisms of somatic hypermutability may differ between distinct patients for a single cancer type [8] and even between distinct cell lineages [9] or over time in a single tumor [10]. Different cancers may be prone to varying degrees of point mutation hypermutability, microsatellite instability, or chromosome instability [11]. Even within these broad classes, there are now numerous recognized “mutator phenotypes” presumed to be caused by distinct hypermutability mutations. For example, approximately thirty point mutation signatures [12, 13] are known to exhibit variability in different cancers, with several either known to be connected to specific kinds of hypermutability defects (e.g., pol-ϵ defect [14], APOBEC defect [15], or various DNA mismatch repair defects such as those are induced by germline *BRCA1* or *BRCA2* mutations [16]), as well as distinct signatures of copy number or structural variation mechanisms [17, 18], such as those due to *TP53* dysfunction [19, 20]. At present, a number of these hypermutability signatures remain of unknown origin [12] and it remains elusive whether others might be detected as we gain better power to resolve broader classes of mutations and precisely quantify them via deep sequencing [21]. A variety of lines of evidence have suggested that these distinct hypermutability phenotypes have important implications for how a tumor is likely to evolve in the future. For example, it has been shown that tumors prone to copy number alterations (CNAs) and aneuploidy via whole genome duplication (WGD) have significantly worse prognoses than similar tumors only prone to focal CNAs [22, 23, 24]. Similar observations have appeared anecdotally for a variety of specific mutation classes.

Here, we sought to explore a key implication of these past studies: how a tumor is likely to progress in the future is influenced by, and in principle predictable from, the mechanisms by which it has evolved so far, independent of the specific spectrum of driver mutations those mechanisms have so far produced. That is, the patient-specific spectrum of mutator phenotypes acting in a given tumor has predictive power towards its future progression. For example, evolutionary statistics based on fluorescence *in situ* hybridization (FISH) data [25] are predictive of whether a tumor will go on to metastasize [26], with more nuanced models of mechanism and variation rate, leading to enhanced predictive power [27, 28]. Conceptually, the use of mutational mechanisms as predictors is distinct from, and complementary to, the standard “driver gene” model of prediction – that we predict likelihoods of tumor progression based on its specific pattern of mutations or expression changes in genes of known functional significance in cancer [29, 30] – which is the basis of much of the current work in genomic diagnostics for cancer. While conventional genomic predictors focus on markers of the selection component of clonal evolution using mutations with putative functional effects on clonal fitness, we propose instead to focus on the diversification component of evolution by profiling phenotypes that affect the degree and kind of mutations a tumor is prone to generate. Here, we specifically develop this idea of evolutionary predictors of progression by applying “tumor phylogenetics”, i.e., the reconstruction of the cell lineage history in a tumor, to infer likely clonal phylogenies of individual tumors and then derive quantitative estimates of the evolutionary processes acting on that tumor from these phylogenies. The combination of mechanisms of mutation and the degrees to which they act in a given cancer should then, we propose, have independent predictive power from its specific driver mutations for future progression. We can harness such predictive power via machine learning frameworks.

The remainder of this paper is devoted to implementing and demonstrating a realization of this idea of progression prediction from somatic hypermutability phenotypes. Below, we describe a general model for this framework with specific variations extracted from whole exome sequence (WES) data and whole genome sequence (WGS) data using a combination of tumor phylogeny methods to extract features for use in predicting progression through regularized Cox regression. Specifically, we demonstrate the effectiveness of this strategy via prediction of overall survival (OS) and recurrence/disease-free survival (DFS) on data from the Cancer Genome Atlas (TCGA) project [31] and the International Cancer Genome Consortium (ICGC) [32], including breast invasive carcinoma (BRCA) [8], lung adenocarcinoma (LUAD) [33], and lung squamous cell carcinoma (LUSC) [34]. In each case, we show that predictions using phylogeny-derived mutator phenotype profiles have significant predictive power for progression outcomes and that incorporating phylogeny-derived evolutionary features leads to enhanced predictive power relative to predictions from clinical and/or traditional driver-centric genomic features alone.

## 2 Results

### 2.1 Overall workflow

Figure 1 summarizes the overall process of variant calling, evolutionary modeling, and regression applied in the present work. We assume that the process begins with tumor genome sequencing data, which include either WES or WGS, and could be either single-sample (solely tumor data), paired tumor/normal, or multi-sample (multiple distinct tumor sites or regions as well as possibly paired normal). While some study designs might alternatively use targeted deep sequencing data, we would generally consider those data not suited to the present methods, which benefit from profiling larger fractions of the genome to estimate better aggregate mutation rates. We consider here only inference from bulk tumor data, although we note that the strategy might be applied to single cell sequence data [35] or combinations of bulk and single-cell data [36, 37], should such data become available for sufficiently large cohorts. The genomic data is preprocessed and passed to one or more variant callers, ideally including single nucleotide variations (SNVs) and copy number alterations (CNAs) calls as well as calls for diverse classes of structural variations (SVs) to produce a variant call format (VCF) file with detected variants and their variant allele frequencies (VAFs) per sample. These variant calls are then passed as input to tumor evolution algorithms, which are used to deconvolve aggregate bulk data into multiple clonal evolutionary states and infer a cellular lineage tree connecting those states and predicting their likely ancestry. Next, a variety of quantitative measures of the evolutionary process, i.e., **evolutionary features**, corresponding to distinct mutation mechanisms are extracted. These features are intended to approximate the degree to which distinct **mutator phenotypes** are activated in a tumor. We evaluate these features either alone or in combination with additional clinical features, such as patient demographics or immunohistochemical profiles, and driver-centric genomic features, which describe the presence or absence of mutations or CNAs in known cancer driver genes. Feature filtering and selection are then conducted and followed by training or applying a regression algorithm for prediction of a clinical outcome of interest.

**Figure 1:**
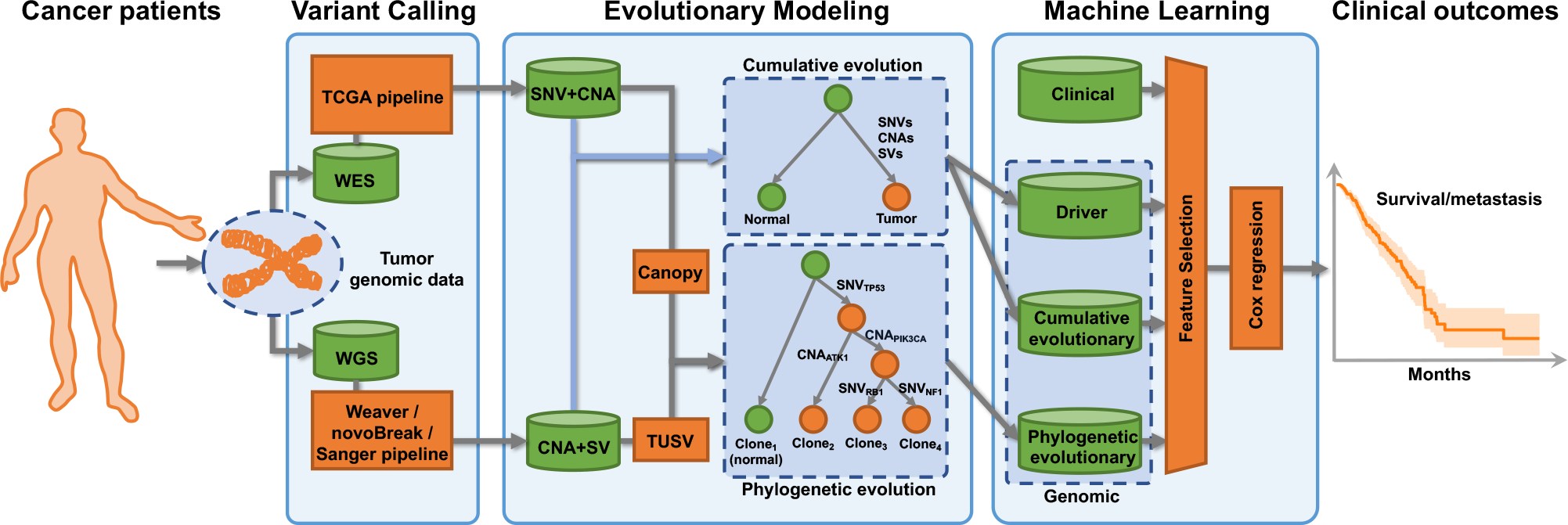
Summary of the prognostic prediction workflow. The overall framework utilizes either WES data followed by standard TCGA variant caller or WGS data followed by either Weaver, novoBreak, or Sanger variant callers to derive measures of mutational preferences from phylogenetic models of clonal evolution and cumulative mutation burdens. The extracted cumulative and phylogenetic evolutionary features, together with clinical metadata and driver features, are used for training and prediction prognoses of patients such as survival and recurrence.

In order to validate the effectiveness and generality of this approach, we compiled three sets of data testing various conditions under which the approach might be applied. These datasets cover:
- Datasets from two data sources: TCGA [31], ICGC [32, 38];
- Three cancer types: BRCA [8], subtypes of lung cancer (LUCA): LUAD [33] + LUSC [34];
- Two different sequencing strategies: WES, WGS;
- Four variant callers: TCGA pipeline [31, 39, 40, 41], Weaver [42, 43], novoBreak [44], Sanger pipeline [45];
- Two phylogenetic methods: Canopy [46] (SNV+CNA), TUSV [47] (CNA+SV);
- Two prognostic prediction tasks: OS, DFS.

We selected breast and lung cancers for validation primarily due to their relatively large TCGA and ICGC cohorts. The details of statistics and settings for these datasets are shown in Table 1.

**Table 1:** Statistics of the experiments and datasets in the study. Three sets of data covering three cancer types (BRCA, LUCA=LUAD+LUSC), two experiment strategies (WES, WGS), four variant callers (TCGA pipeline, Weaver, novoBreak, Sanger pipeline), two phylogenetic models, and two prediction tasks (OS, DFS) were conducted. We omit the DFS prediction task for LUAD cancer in Exp 2 as there were only three positive examples.

| Exp | Dataset | Cancer | Seq | Variant | Phylogenetic | Size | Event/Censored |  |
| --- | --- | --- | --- | --- | --- | --- | --- | --- |
|  |  | Type | Strategy | Caller | Model |  | OS | DFS |
| 1 | TCGA | BRCA | WES | TCGA | Canopy | 1044 | 145/897 | 102/764 |
|  |  | LUAD | WES | TCGA | Canopy | 516 | 182/325 | 181/268 |
| 2 | TCGA | BRCA | WGS | Weaver | TUSV | 28 | 5/23 | 3/19 |
|  |  | LUAD | WGS | novoBreak | TUSV | 59 | 29/30 | 25/18 |
| 3 | ICGC | BRCA | WGS | Sanger | TUSV | 90 | 17/73 | 14/59 |
|  |  | LUCA | WGS | Sanger | TUSV | 89 | 44/43 | 29/38 |

### 2.2 Genomic features are complementary to clinical features

We first explored the overall correlation structure of the full feature space in order to identify the orthogonal features and provide biological insight into subgroups of features, as well as to validate the min-redundancy rule used by our feature selection method (Sec. 5.3). We refer the reader to Sec. 5.2 for full lists of features considered in each class. We calculated the Pearson correlation coefficient between each pair of features filtered by the max-relevance rule, and performed hierarchical clustering based on Ward distance [48] to group them. Since Exp 1 has many more samples compared to Exp 2 and 3 and thus provides the most reliable and robust results, we mainly focus here on Exp 1.

Figure 2 shows the analysis on the features of the BRCA OS task in Exp 1. The figure allows one to distinguish three or four closely related blocks of features along the diagonal, corresponding to generic evolutionary features (block-e), clinical features (block-c), cumulative evolutionary features (block-u), and driver features (block-d). **Block-e:** Most of the phylogenetic and cumulative evolutionary features not related to CNAs are collapsed into a single high-correlation block essentially corresponding to point mutation rates. This correlation can be easily understood, since all of the features in the block capture overall SNV rate in various ways, although phylogenetic features are more fine-grained than cumulative features. For example, the SNV rates in the largest clone (*lg clone snv*; a phylogenetic evolutionary feature) should be close to the total SNV rates (*snv rate*; a cumulative evolutionary feature) and therefore they should be positively correlated. **Block-c:** Most clinical features are negatively, rather than positively, correlated with other clinical features outside of small, highly correlated sub-blocks. The main reason comes from the data processing and feature extraction step when we map categorical clinical features into one-hot vectors and therefore introduce collinearity (Sec. 5.2). For example, the *breast carcinoma progesterone receptor status* | *positive* and *breast carcinoma progesterone receptor status* | *negative* originally came from the same categorical clinical feature *breast carcinoma progesterone receptor status* to yield two anti-correlated binary features. We can easily break this type of collinearity by dropping one of the anti-correlated categories. Strong positive correlation is also sometimes observed within clinical features for biological reasons, such as that between PR status (*breast carcinoma progesterone receptor status* | *positive*) and ER status (*breast carcinoma estrogen receptor status* | *positive*) reflecting a common association with luminal breast cancer subtypes [49], or that between *age at initial pathological diagnosis* and *menopause status*. **Block-u**: This block essentially collects CNA rate features, which we note are highly correlated with one another but only weakly correlated with generic tree topology features that are influenced by both SNV and CNA rates. **Block-d:** Driver features largely form their own distinct block of high mutual correlation. This block is moderately correlated with both SNV and CNA rate features. We hypothesize that the generic pattern of positive correlation among driver features reflects generic differences in background SNV or CNA rates from different somatic hypermutability phenotypes [2]. For example, we propose that tumors exhibiting one CNA driver mutation are more likely to exhibit other CNA driver mutations because they are more likely to have an elevated rate of CNA mutations generically. This interpretation is confirmed by the cross-correlations between driver features and CNA rate features (block-ud) and between driver and SNV rate features (block-ed).

**Figure 2:**
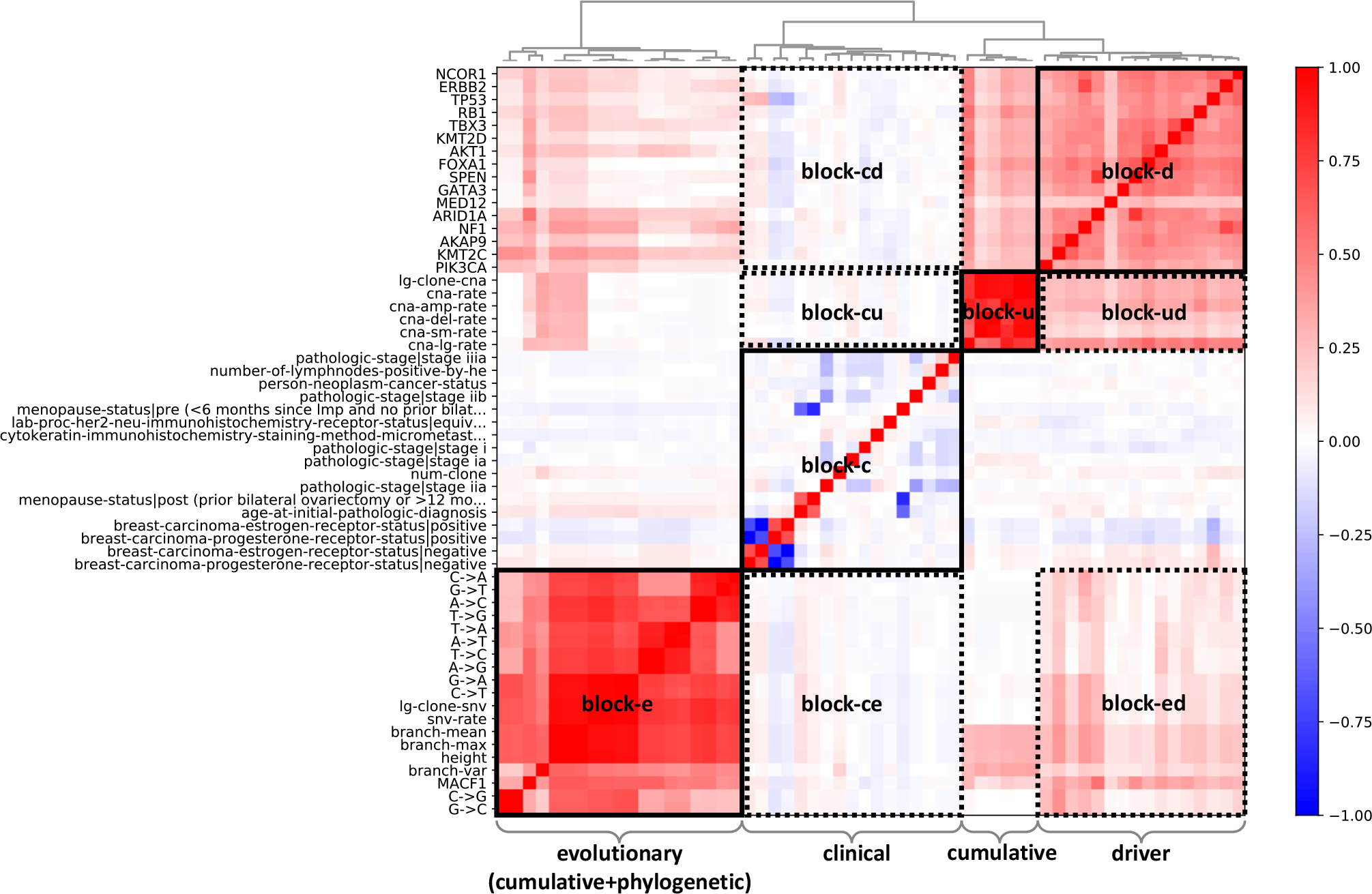
Heatmap of correlation for clinical, driver, and evolutionary features. Pearson correlations of features filtered by max-relevance rule for BRCA OS task in Exp 1 are calculated. Therefore, each shown feature can be used for effectively predicting OS in BRCA samples. Due to display limit, we only show the feature name along each row, while the features of columns are in the same order as rows. Strong correlation within each feature type is observed (block-e, block-c, block-u, block-d), while genomic features are more independent and orthogonal to clinical features (block-cd, block-cu, block-ce).

In general, the features from each of the four main feature blocks are highly correlated to other features of the same block, validating that the min-redundancy rule is necessary to prevent collinearity and reduce model complexity. On the other hand, the blocks are relatively independent of each other aside from the previously noted generic correlations between driver features and mutational rates. This independence is especially true for genomic features generically vs. clinical features (white block-cd, block-cu, and block-ce). Therefore, one may conclude that the genomic features are roughly orthogonal to the clinical features and can be used as complementary in the task of prognostic prediction to possibly improve the performance. We also note that our analysis are concordant with findings of Dawson et al. [50] that suggested a partitioning of breast cancers in part into distinct classes of SNV-driven and CNA-driven tumors, although our findings are consistent with the observation that these are two orthogonal classes of features that may act to different degrees in all tumors as opposed to orthogonal classes of tumors.

We show the correlation results of all the three experiments, two tasks and two cancer types in the Supplementary Materials (Fig. S1-S10). In general, similar correlation patterns mentioned above still exist, although in some cases, the cumulative evolutionary features will be closer to phylogenetic evolutionary features other than driver ones. One may also observe that SV-related features that are not available for the WES experiments but are visible in the WGS experiments (Exp 2-3; Fig. S4-S10), such as total SV rates (*sv rate*), are positively correlated with total CNA rates (*cna rate*), likely reflecting the fact that CNAs are typically interwoven with SVs of various kinds. In addition, the SV-related features are also positively correlated with phylogenetic evolutionary features such as the height of the phylogeny (*height*), which are influenced by all forms of mutation processes.

### 2.3 Genomic features improve prognostic prediction

It is expected that clinical features will have the strongest individual predictive value and we are primarily interested here in establishing whether evolutionary features enhance the predictive value of clinical outcomes relative to clinical features alone or clinical features supplemented by driver genomic features. Our three groups of evaluation datasets were intended to explore different settings and possibilities in clinical practice (Table 1), such as different cancer types, sequencing methods, and prediction tasks, to answer that question. The results indicate that genomic features of both classes do indeed enhance the predictive power (Table 2, 3, Fig. 4) and that evolutionary features enhance predictive power relative to clinical, driver, or the combination of the two feature classes.

**Table 2:** Performance of prognostic prediction with different features in Exp 1 WES samples. Performance is evaluated by the concordance index (CI; Sec. 5.4) on the test set, with results showing mean ± standard deviation for each test. Experiments were implemented using 10-fold CV, and repeated for five times to calculate the means and standard deviations to measure the robustness of the model. “genomic” means all genomic features, including driver, cumulative evolutionary and phyloge-netic evolutionary features. “clinicalΔ” means the neoplasm status (*person neoplasm cancer status*) is removed from clinical features. Clinical feature set performs best among all the four feature sets. However, the additional genomic features are always synergistic and promote the prediction of prognoses. Statistical significance notation for the “clinical+driver/cumulative/phylogenetic/genomic” vs. “clinical”, and “clinicalΔ+driver/cumulative/phylogenetic/genomic” vs. “clinicalΔ” is defined by the one-sided test *p*-value. *: *p*< 0.05; **: *p* < 0.01; ***: *p* < 0.001.

| Feature | Experiment 1 |  |  |  |
| --- | --- | --- | --- | --- |
|  | BRCA |  | LUAD |  |
|  | OS | DFS | OS | DFS |
| clinical | 82.0±0.21 | 78.4±0.42 | 69.3±0.31 | 65.5±0.33 |
| driver | 57.8±0.60 | 57.0±1.14 | 53.2±1.12 | 53.8±1.16 |
| cumulative | 58.8±1.57 | 53.7±1.05 | 52.3±0.49 | 55.4±0.47 |
| phylogenetic | 56.9±1.06 | 52.0±0.63 | 51.2±0.25 | 52.8±0.36 |
| clinical+driver | 82.4±0.26** | 79.0±0.21*** | 69.6±0.21* | 66.2±0.33** |
| clinical+cumulative | 82.1±0.24 | 78.4±0.43 | 69.7±0.50 | 67.0±0.21*** |
| clinical+phylogenetic | 82.0±0.21 | 78.5±0.43 | 69.5±0.31 | 66.1±0.16*** |
| clinical+genomic | **82.5±0.24\*\*\*** | **79.1±0.27\*\*\*** | **69.9±0.25\*\*\*** | **67.7±0.18\*\*\*** |
| clinicalΔ | 74.6±0.33 | 70.3±0.41 | 65.5±0.23 | 60.3±0.40 |
| clinicalΔ+driver | 75.0±0.22** | 71.3±0.70** | 66.9±0.41*** | 61.9±0.57*** |
| clinicalΔ+cumulative | 74.8±0.40 | 70.5±0.45 | 66.7±0.24*** | 61.6±0.35*** |
| clinicalΔ+phylogenetic | 74.7±0.37 | 70.3±0.41 | 66.4±0.29*** | 61.4±0.37*** |
| clinicalΔ+genomic | **75.1±0.24\*\*** | **71.7±0.57\*\*\*** | **67.4±0.30\*\*\*** | **63.1±0.69\*\*\*** |

**Table 3:** Performance of prognostic prediction (CI) with different features in Exp 2 and Exp 3 WGS samples. All the results were based on LOOCV. The DFS task of BRCA in Exp 2 was not conducted due to limited positive samples (three). Similar to the results in Exp 1 (Table 2), the additional genomic features always yield improved prediction performance.

| Feature | Experiment 2 |  |  |  | Experiment 3 |  |  |  |
| --- | --- | --- | --- | --- | --- | --- | --- | --- |
|  | BRCA |  | LUAD |  | BRCA |  | LUCA |  |
|  | OS | DFS | OS | DFS | OS | DFS | OS | DFS |
| clinical | 80.2 | - | 64.3 | 56.3 | 73.8 | 84.8 | 69.4 | 67.5 |
| driver | 75.3 | - | 33.0 | 31.7 | 66.0 | 76.6 | 60.7 | 55.2 |
| cumulative | 60.5 | - | 55.2 | 59.1 | 54.4 | 56.5 | 57.3 | 57.5 |
| phylogenetic | 61.7 | - | 57.3 | 51.7 | 54.6 | 59.0 | 43.6 | 44.2 |
| clinical+driver | **85.2** | - | 64.8 | 56.5 | 76.8 | 87.7 | 71.3 | 71.7 |
| clinical+cumulative | 80.2 | - | 68.2 | 65.0 | 73.8 | 87.3 | **72.1** | 74.5 |
| clinical+phylogenetic | 80.2 | - | 67.1 | 62.5 | 74.0 | 85.0 | 69.4 | 67.8 |
| clinical+genomic | **85.2** | - | **70.0** | **66.4** | **77.7** | **91.1** | **72.1** | **76.5** |
| clinicalΔ | 55.6 | - | 58.0 | 48.7 | 61.0 | 71.8 | 64.6 | 62.5 |
| clinicalΔ+driver | **71.6** | - | 62.3 | 48.7 | **71.1** | 79.3 | **68.1** | 68.9 |
| clinicalΔ+cumulative | 56.8 | - | 70.1 | 54.3 | 69.1 | 72.5 | **68.1** | 75.0 |
| clinicalΔ+phylogenetic | 69.1 | - | 69.9 | 53.6 | 69.1 | 71.8 | 65.7 | 65.4 |
| clinicalΔ+genomic | **71.6** | - | **71.7** | **57.3** | **71.1** | **81.1** | **68.1** | **76.6** |

Table 2 summarizes OS and DFS prediction performance in Exp 1 across feature classes and combinations of classes in comparison to clinical predictors alone. The performance is evaluated in the metric of concordance index (CI; Sec. 5.4) on the test set. Exp 1 uses WES data and contains the largest corpus of samples available (1044 for BRCA and 516 for LUAD). We utilized 10-fold cross-validation (CV) to train and evaluate. In order to quantify the uncertainty resulting from random choices in splitting the datasets and assess the robustness of the methods, the experiments were repeated five times to derive the means and standard deviations of prediction outcomes. We hypothesize that clinical predictors should provide the strongest predictive information among any single predictive class and do indeed find that to be the case, as assessed by CI. We find that for OS task on BRCA WES data, clinical features outperform driver and cumulative evolutionary prediction, but either clinical+driver or clinical+cumulative features tend to slightly outperform clinical alone. Similarly, for the DFS task on BRCA WES data, either clinical+driver or clinical+phylogenetic combinations tend to perform slightly better than clinical alone. In both OS and DFS tasks for LUAD WES data, clinical prediction outperforms driver, cumulative or phylogenetic prediction, but clinical plus any one of the genomic feature classes again tends to slightly outperform clinical features alone. We further compared to the combination of clinical and all three genomic feature classes (denoted as “clinical+genomic”). In general, the improvements achieved by introducing genomic features are slightly larger for LUAD than for BRCA. This might be due to a relatively more comprehensive clinical description for BRCA than LUAD, making it harder to achieve improvements over clinical features in BRCA. In all cases for Exp 1, clinical+genomic features proves to be the strongest combination, albeit by generally small amounts when inferring from WES data alone, suggesting that each feature class adds independent predictive information.

The clinical feature neoplasm status (*person neoplasm cancer status*), which indicates whether the patient is tumor-free or with tumor, is a strong clinical feature that significantly improves the performance of our model. Both breast and lung cancer patients have distinct survival and recurrence probabilities for different neoplasm status (Fig. 3) and it is the single strongest predictor of outcome, but its availability could be considered unreasonable to assume in normal practice. We were therefore interested in how the models behave without it. For the clinical feature set in the absence of that feature (dubbed “clinicalΔ”), we find a qualitatively similar result that clinical prediction value is still superior to driver or evolutionary alone, although to a lesser degree, but the clinicalΔ+genomic feature combination still outperforms clinicalΔ alone in both cancer types and prediction tasks. In addition, the improvements from the additional genomic features in the case when neoplasm status is missing are in general larger than the case with the full clinical feature set. The improvements of clinical+genomic prediction relative to clinical alone are significant in every case of Exp 1, using a one-sided statistical test: *p*-value < 0.001 for most cases; *p*-value < 0.01 for “clinicalΔ+genomic” vs. “clinicalΔ” in the BRCA OS task.

**Figure 3:**
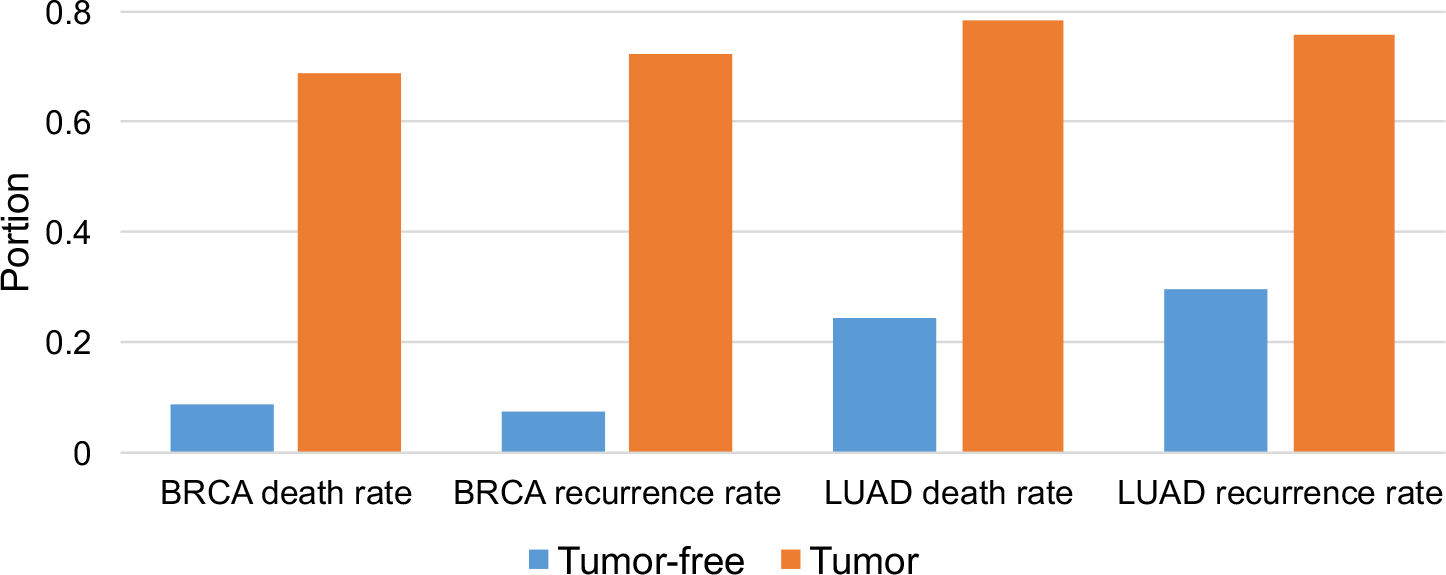
Conditional distribution of death or recurrence rates given the neoplasm status clinical feature in both BRCA and LUAD samples. Patients with positive neoplasm status are much more prone to death or metastasis compared to tumor-free patients, indicating that neoplasm status (*person neoplasm cancer status*) is a strong covariate for our regression model. The distribution is plotted using samples in Exp 1.

Comparable analyses for Exp 2 and Exp 3, which are based on much smaller cohorts but with WGS data allowing for SV discovery, yielded qualitatively similar observations to the WES experiments but with generally substantially larger absolute improvements in accuracy from incorporating evolutionary features in the predictions. We omitted one experiment (BRCA DFS prediction in Exp 2) because the cohort contained only three positive examples, which was too few to allow for cross-validated training. The results are shown in Table 3. Leave-one-out CV (LOOCV) was implemented to train and evaluate the performance, although the cohorts were too small to assess the statistical significance of improvements. Similar observations can be found in these two experimental settings to those from Exp 1. The full clinical feature class is always the single strongest feature set, while the additional genomic features can always improve on these and achieve the best performance among all combinations of feature sets. In most cases, clinical+genomic features improve upon all combinations of subsets of features, although a few instances show ties between clinical+genomic and clinical+driver or clinical+cumulative. We also observe that specific genomic features may not be predictive in some cases. For example, the driver features of LUAD in Exp 2 and phylogenetic features of LUCA in Exp 3 do not work well for predicting OS or DFS. This variability may come from the different variant calling pipelines used by the different sequencing projects and indicates the importance of feature engineering for the prognostic prediction task. One may compare the relative improvements that benefit from genomic features across all the three experiments. It can be observed that the improvements derived from including genomic features are larger for the WGS experiments (Exp 2, 3) than for the WES experiment (Exp 1). This may indicate that a sequencing method that covers a larger area of the genome is important for providing effective and informative genomic features, that the SV features available only in the WGS experiments add useful orthogonal predictive power, or that there is greater synergistic value in the combined features for reducing uncertainty when cohort sizes are small.

Finally, we evaluated the ability of the best combinations of feature classes in Exp 1-3 to stratify patients with distinct censored survival/non-recurrence time using the logrank test (Fig. 4). Patients were classified into two groups: malignant and benign, based on the predicted hazards of the events (decease or recurrence). Samples with predicted hazards larger than the median of predictions were classified as malignant, otherwise as benign. Figure 4a illustrates the separations by OS and DFS time between the predicted malignant and predicted benign WES tumors in Exp 1 through distinct Kaplan-Meier estimator curves, which are significant in both breast and lung cancers quantitatively measured by *p*-values. Similar significant separations exist for WGS tumors in Exp 2 and 3 (Fig. 4b,c). The only exception – the DFS task of LUAD in Exp 2 is not significant (*p*-value > 0.05) – may be explained by the small number of WGS samples in the TCGA LUAD cohort.

**Figure 4:**
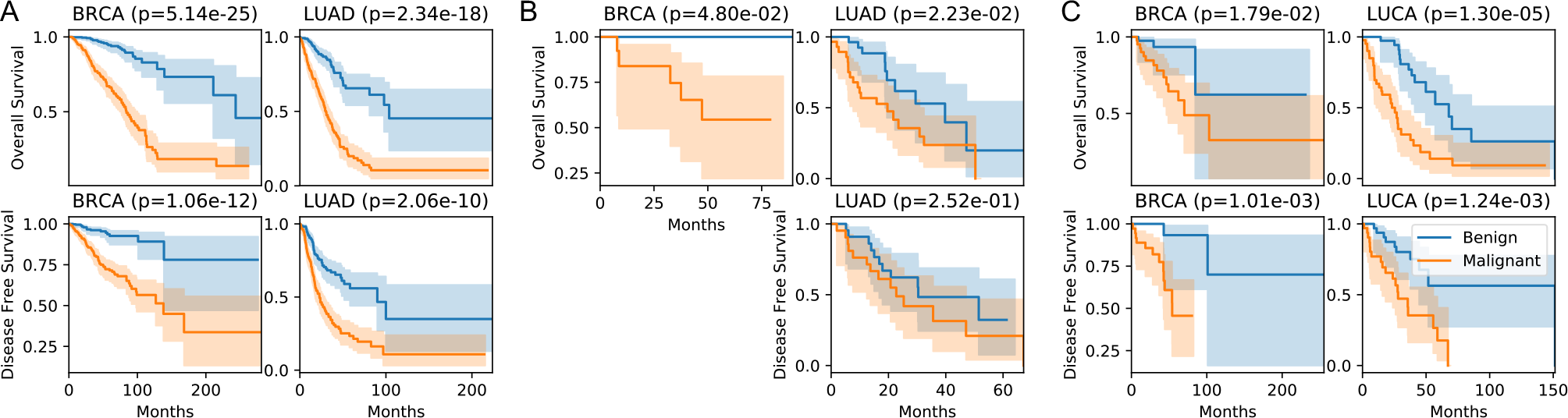
Kaplan-Meier estimators of predicted malignant and benign samples using both clinical and genomic features in (a) Exp 1, (b) Exp 2, and (c) Exp 3. Logrank test shows significant distinct survival and recurrence profiles in both two prediction tasks and two cancer types. The separation of LUAD DFS task in Exp 2 is not significant (*p*-value > 0.05), partly due to the limited number of samples. No experiments were done on the OS task of BRCA in Exp 2 due to the limited number of available positive samples (three).

In summary, these experiments indicate that genomic and clinical features act synergistically to improve the prediction of prognoses, and that in general adding evolutionary features that capture mutational preferences of tumors enhances predictive power relative to clinical features, driver features, or their combination.

### 2.4 Informative features for predicting clinical outcomes

We next sought to assess the contributions of individual features and feature subgroups to the predictive power. We assessed whether an individual feature is important to prognostic prediction in two ways. The first way is to assess the predictive power of the single feature for the task, i.e., univariate regression analysis, and then aggregate them. The second way is to find the set of features that together achieve the best performance, e.g., multivariate regression analysis. These correspond to the two stages of feature selection employed by our model selection strategy (Sec. 5.3): max-relevance filtering and step-wise selection. We therefore analyze and compare these two lists of features.

We mainly focus here on the features selected in Exp 1, since they derive from much larger cohort sizes and are therefore are more statistically reliable. The clinical, driver, cumulative evolutionary, and phylogenetic evolutionary features that are predictive (CI > 0.5) for the four cases (OS or DFS in breast or lung cancers) are sorted in Fig. 5. **Universal features across tasks and cancer types:** As one can see, the top clinical feature for both breast and lung cancers is neoplasm status (*person neoplasm cancer status*), which reports whether the patient has a tumor or is tumor-free. The prognoses are significantly more optimistic when no tumor is observed in general (Fig. 3). In addition, *pathologic stage* is the second universal predictive clinical feature, which is also unsurprising and reflects the known value of expert staging in predicting cancer outcome [51]. The most strongly predictive evolutionary features vary by tumor type and task, with both SNV and CNA features showing up as significant. Cumulative evolutionary feature of total CNA rates (*cnv rate*), including rates of CNA above/below 500,000 nt (*cnv lg rate*, *cnv sm rate*) and CNA duplication/deletion rates (*cnv amp rate*, *cnv del rate*), emerge as important in all four cases, indicating that the CNA rates and various sub-categories of it are broadly important predictors of progression. Phylogenetic evolutionary features such as the variance of edge lengths (*branch var*; predictive in all cases except BRCA DFS), and CNA rates in the largest clone (*lg clone cnv*) are shared informative single features as well. These observations are consistent with the notion that evolutionary processes that promote high heterogeneity are predictive of poor outcome generically [52]. **Informative features of specific cancer type:** We also observe that the informative features for OS and DFS prediction tasks are extensively shared within the same cancer type. For example, our finding that the clinical features PR status (*breast carcinoma progesterone receptor status*) and ER status (*breast carcinoma estrogen receptor status*) are highly informative for both tasks is consistent with the current subtype classification method of breast cancer based on hormone receptor status [49]. The clinical features *number of lymphnodes positive by he*, *cytokeratin immunohistochemistry staining method micrometastasis indicator* are consistent with previous research on the value of these factors as predictors of breast cancer survival [53] and metastasis [54]. While in the present work, we directly take cytokeratin immunohistochemistry staining as a feature, one should note the possible artifacts that could be introduced in practice [55]. A large number of driver features, including *KMT2C*, *KMT2D*, *FOXA1*, *TBX3*, *AKAP9*, *MED12*, *GATA3*, and *AKT1* are informative for both OS and DFS prediction in breast cancer [29, 56]. Cumulative evolutionary features *C* → *T*, *T* → *C*, *G* → *A*, *G* → *C* are informative for both OS and DFS prediction in breast cancer. The SNV rate of *C* → *T* is especially important for the breast cancer prognostic prediction. *C* → *T* (and, equivalently, *G* → *A*) preferences are associated with several mutational signatures [12] and we would hypothesize that their association with age-related signatures largely accounts for their predictive power here (Fig. 2). The relevance of *G* → *C* (equivalently, *C* → *G*) may reflect its association with APOBEC dysfunction [15]. For the BRCA OS specifically, we find the cumulative evolutionary feature of total SNV rates (*snv rate*) and the phylogenetic evolutionary feature SNV rates in the largest clone (*lg clone snv*) are predictive [2]. In addition, phylogenetic evolutionary features such as the height of the phylogeny (*height*), average of branch lengths (*branch mean*), and maximum edge length (*branch max*) are informative for both OS and DFS of BRCA, suggesting again that overall measures of evolutionary heterogeneity are important to breast cancer progression and prognoses [26]. In lung cancer, the clinical features *anatomic neoplasm subdivision* and *histological type* are predictive for both OS and DFS, while *gender* is predictive for OS, consistent with the previous research that the right lung and male gender are usually associated with worse prognoses [57]. The driver feature *LRP1B* is informative for both prediction tasks in lung cancer [58]. **Informative features distinct between OS and DFS tasks:** Although the informative features mainly differ across cancer types, we did note that the clinical feature *age at initial pathologic diagnosis* is only informative for predicting OS and not DFS, in both BRCA and LUAD samples. This may reflect the fact that older patients are more likely to die of other competing risks, such as heart attack, than their cancer during the time of follow-up [59], although it could potentially also involve cancer-intrinsic effects of somatic mutation processes likely having been active longer in tumors of older versus younger patients. Teasing apart the various confounding factors introduced by age at diagnosis is a complex question, however, beyond the scope of the present study [60].

**Figure 5:**
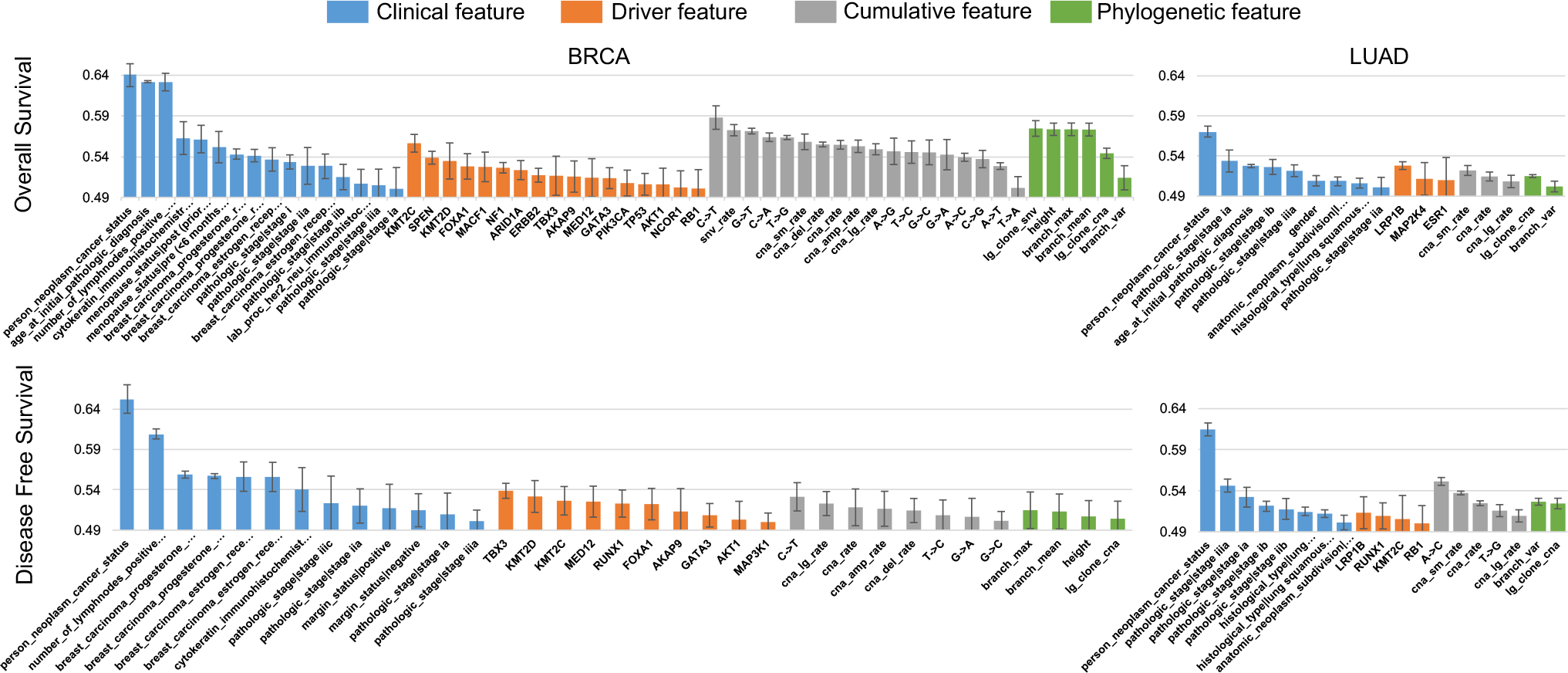
List of single predictive features of prognoses. The features are sorted in the descending order of their prognostic prediction performance for BRCA and LUAD in Exp 1. The features whose CI larger than 0.5 using univariate Cox regression are shown.

Apart from the single informative features, the combination of features that lead to the best performance in Exp 1 are collected in Table 4, i.e., the selected clinical and genomic features that have the optimal performance in clinical+driver, clinical+cumulative, and clinical+phylogenetic experiments. The selected clinical features are roughly the same set as the single informative clinical features, as we might expect given that clinical features generally show a fairly low correlation with one another. However, not every individual informative feature is finally selected due to redundancy between features. Even well-established clinical features, such as ER status, are not reliably selected by the final best predictor, presumably because the information they carry is redundant with that derivable from the other features. These final sets of selected features are different from the ones filtered by the max-relevance rule, in that redundant features are removed to produce a collectively non-redundant set. Therefore the final selected clinical features form a subset of the filtered features. For genomic data, the method selects not just individually informative features, but also additional not-so-predictive features that prove predictive in combination with others. The *PIK3CA*, *MACF1* and *KMT2C* features for BRCA OS; *MED12* for BRCA DFS; *LRP1B* for LUAD OS; *LRP1B* and *RB1* for LUAD DFS — all of which are among the single informative drivers — are selected finally as complementary features. However, the additional driver feature *CDH1* is selected for the DFS prediction task of LUAD despite not showing up as significantly predictive individual [61], indicating that a feature that is not able to predict accurately by itself may help to improve a prediction model when combined with other features. For evolutionary features, only a subset of the individual predictive features are selected finally, such as *G* → *T* (in BRCA OS), rates of CNA below 500,000 nt (*cna sm rate*; in LUAD OS), *A* → *C* (in LUAD DFS), CNA rates in the largest clone (*lg clone cna*; in LUAD OS and DFS). This is to be expected given the correlations previously observed between subclasses of evolutionary features. We further note that it is likely these best feature sets are not unique and we would expect that there may be distinct combinations of features that collectively perform as well or nearly so as the set considered here.

**Table 4:** Finally selected features for predicting prognoses in Exp 1.

| Cancer | Feature | Task |  |
| --- | --- | --- | --- |
|  |  | OS | DFS |
| BRCA | Clinical | person neoplasm cancer status<br>age at initial pathologic diagnosis<br>pathologic stage<br>breast carcinoma progesterone receptor status<br>menopause status<br>cytokeratin immunohistochemistry staining method<br>micrometastasis indicator | person neoplasm cancer status<br>breast carcinoma progesterone receptor status<br>number of lymphnodes positive by he<br>pathologic stage<br>margin status<br>cytokeratin immunohistochemistry staining method<br>micrometastasis indicator<br>breast carcinoma estrogen receptor status |
|  | Driver | *PIK3CA*<br>*MACF1*<br>*KMT2C* | *MED12* |
| | Cumulative | $G \rightarrow T$ | ∅ |
| | Phylogenetic | ∅ | ∅ |
| LUAD | Clinical | person neoplasm cancer status<br>pathologic stage<br>age at initial pathologic diagnosis | person neoplasm cancer status<br>pathologic stage<br>histological type |
|  | Driver | *LRP1B* | *LRP1B*<br>*RB1*<br>*CDH1* |
| | Cumulative | rates of CNA below 500,000 nt | $A \rightarrow C$ |
|  | Phylogenetic | CNA rates in the largest clone | CNA rates in the largest clone |

The finally selected features for Exp 2 and Exp 3 are shown in Table 5, 6. They provide results using WGS data in contrast to the WES data in Exp 1. We compare their final results with that of Exp 1 in Table 4. One can see that the selected clinical features appear to be consistent across three groups of experiments. The selected driver features are more variable. Additional driver features can improve the corresponding performance, such as *ESR1* (Exp 2 LUAD OS), *AKAP9* (Exp 3 BRCA OS), *ESR1* (Exp 3 BRCA DFS), and *SVEP1* (Exp 3 LUCA DFS), although *ESR1* and *AKAP9* are still among the list of single informative features and not finally selected in Exp 1. The difference may partly come from the LUSC samples, which are not included in Exp 1 and 2. With the additional SV data available, Exp 2 and 3 (WGS) capture similar but not identical sets of evolutionary features. The selected cumulative evolutionary features include *A* → *C*, *G* → *T*, rates of CNA below/above 500,000 nt, similar to that selected in Exp 1 (WES). However, Exp 2 and 3 additionally select some SV-related phylogenetic evolutionary features which could not be derived for Exp 1, e.g., average/variance of edge lengths in the unit of SV rates. This is consistent with prior knowledge that SVs are a crucial mechanism of tumor progression and functional adaptation through their role in creating CNAs as well as fusion genes [62] and contribute substantially to tumor evolution [47].

**Table 5:** Finally selected features for predicting prognoses in Exp 2.

| Cancer | Feature | Task |  |
| --- | --- | --- | --- |
|  |  | OS | DFS |
| BRCA | Clinical | person neoplasm cancer status<br>pathologic stage | - |
|  | Driver | *MACF1* | - |
| | Cumulative | ∅ | - |
| | Phylogenetic | ∅ | - |
| LUAD | Clinical | pathologic stage<br>person neoplasm cancer status<br>anatomic neoplasm subdivision<br>race | person neoplasm cancer status<br>pathologic stage<br>race |
|  | Driver | *ESR1* | *LRP1B* |
| | Cumulative | rates of CNA above 500,000 nt | rates of CNA above 500,000 nt<br>$A \rightarrow C$ |
|  | Phylogenetic | variance of edge lengths | average of edge lengths in unit of SV rates<br>portion of the largest clone |

**Table 6:** Finally selected features for predicting prognoses in Exp 3.

| Cancer | Feature | Task |  |
| --- | --- | --- | --- |
|  |  | OS | DFS |
| BRCA | Clinical | pathologic stage<br>menopause status<br>person neoplasm cancer status<br>breast carcinoma estrogen receptor status | person neoplasm cancer status<br>race<br>cytokeratin immunohistochemistry staining method<br>micrometastasis indicator<br>pathologic stage |
|  | Driver | *MACF1*<br>*AKAP9* | *ESR1* |
| | Cumulative | ∅ | rates of CNA below 500,000 nt |
|  | Phylogenetic | variance of edge lengths in unit of SV rates | variance of edge lengths in unit of CNA rates |
| LUCA | Clinical | pathologic stage<br>anatomic neoplasm subdivision<br>person neoplasm cancer status<br>gender | person neoplasm cancer status<br>pathologic stage<br>gender<br>race |
|  | Driver | *LRP1B* | *SVEP1* |
| | Cumulative | $G \rightarrow T$ | $G \rightarrow T$<br>rates of CNA above 500,000 nt |
| | Phylogenetic | ∅ | average of edge lengths in unit of SV rates |

### 2.5 Landscapes of cancer patients in evolutionary feature space

One of the major observations above is that genomic features partition into subsets with distinctive patterns of correlation within and between them (Sec. 2.2). We sought to explore aspects of the correlation structure that might not be readily apparent solely from the pairwise correlation analysis. To investigate the landscape of tumors in the evolutionary feature space, we considered all the cumulative evolutionary and phylogenetic evolutionary features that passed the filtering step by removing the redundant information or noise. In order to visualize the manifold defined by cancer patients in this space of evolutionary features, we conducted principal component analysis (PCA). We show the first two principal components (PCs) in Fig. 6. Each sample is represented as a grey dot. The survival time in the feature space is interpolated using *k*-nearest neighbors (*k*-NN) on deceased samples to reduce the noise from survival data of individual samples. *k* was chosen to be 40, which is large enough to smooth the estimated OS/DFS, while small enough to keep enough details in the feature space. The contours of the estimated surfaces are then shown as well, where darker color represents shorter estimated survival time or recurrence time and thus more malignant status. We can estimate the contours more reliably with larger sample sizes and therefore focus again here on the data in Exp 1. Note that there exist local artifacts of the contours using *k*-NN, when *k* is larger than one, e.g., the lighter islands in the bottom left of Fig. 6b and top left of Fig. 6d. We map one of the cumulative features and phylogenetic features into the first two PCs as well for interpretation purposes, specifically picking for each plot the two features with largest magnitudes of coefficients in regression.

**Figure 6:**
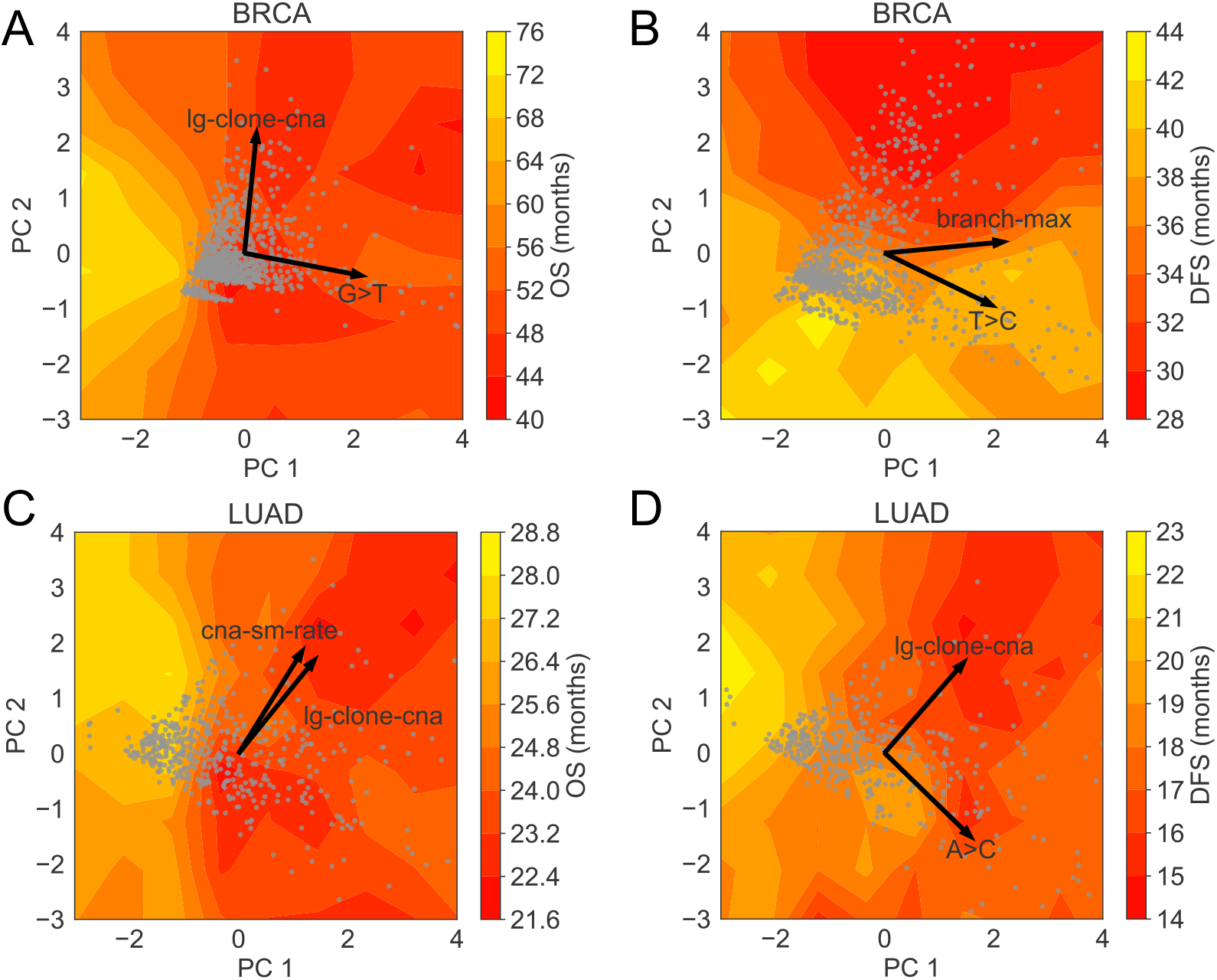
OS and DFS landscapes of BRCA and LUAD cancer patients. Figures are plotted based on samples in Exp 1. PCA was conducted on the samples using the cumulative and phylogenetic evolutionary features that passed the filtering step of feature selection. Each patient is represented as a single grey dot in the figure. The contour of survival time in the two-dimensional manifold was estimated based on the *k*-NN algorithm. One finally selected cumulative and one finally selected phylogenetic features in each case are shown in the PCA plot as well for the case study.

As the figures show, most of the patients roughly locate in a triangular manifold, suggestive of a large number of patients with similar mutation profiles, but with long tails of outliers in each dimension. Consistent with the pairwise analysis, we observe that CNA and SNV features are largely orthogonal (e.g., *C* → *T* vs. CNA rates in the largest clone; *A* → *C* vs. CNA rates in the largest clone) and pairs of features from either class alone are close to parallel (e.g., rates of CNA below 500,000 nt vs. CNA rates in the largest clone), with generic evolutionary features in between. We do not see any obvious partitioning by tumors into two distinct sub-manifolds along the orthogonal axes, again suggesting a subtly different observation from Dawson et al. [50] that CNA and SNV mutation preferences are largely orthogonal factors in the mutational landscape but do not necessarily lead to orthogonal sets of tumors as individual tumors may exhibit variation in both factors independently. There exist patterns and internal structures of OS or DFS time for both BRCA and LUAD patients in the first two PCs. Roughly the patients that lie in one side of the evolutionary feature space (light area) have more optimistic clinical outcomes than the ones in the other half feature space (dark area). The figures show correlation between filtered evolutionary features and OS/DFS although the outcomes are not obviously strongly correlated to any one specific feature.

## 3 Discussion

Our work supports the central hypothesis that variation in mutational phenotypes is predictive of progression outcomes, independent of clinical factors and specific driver gene mutations, but large gaps remain in exploring in detail the space of predictors, the mechanisms by which they act, and the best strategies to realize their translational potential. Some of these questions remain data-limited and likely cannot be answered without larger cohorts and richer clinical metadata. Our work suggests that WGS provides an advantage over WES in improving prediction power beyond that of clinical and driver-centric features, but available cohorts of patients with WGS data are still limited, an issue that proved a limitation for our Exp 2 and Exp 3 despite their use of some of the largest cancer WGS corpora that have been made accessible to researchers. The lack of samples could lead to fragile models and statistically insignificant performance.

Another area in which the work can likely be advanced is to effectively describe patients-specific mutational phenotypes through a set of partially independent variables. Our results suggest that CNA and SNV mutator phenotypes act essentially independently of one another. Prior literature suggests a finer structure underlying these two broad types of process exists. In the SNV domain, mutational signature analysis has demonstrated the existence of approximately 30 independent point mutation processes acting to different degrees on different cancers [12]. We likewise know that distinct mechanisms of SV and CNA act on a tumor genome [63] and at least some of these, such as whole genome duplication (WGD), are independently predictive of outcome [22, 23, 24]. Our PCA analysis of the evolutionary feature space did not reveal strong structure that would directly allow us to reconstruct the intrinsic space of orthogonal mutability features, which may reflect the high variability of the mutational processes, the limits of the sample size, the imprecision of rate estimates, and the limitations of PCA in reconstructing what may be a non-linear low-dimensional manifold describing the true space of hypermutability processes acting in cancers. Additional efforts may need to be paid in discovering the more consistent and complicated manifolds of samples in both BRCA or LUCA.

There are many avenues by which the predictive power of the model might also be optimized. First, although the current evolutionary features provide improvements in addition to the clinical and driver features, we may boost more by integrating the results from different variant calling pipelines. The technique of ensemble learning and boosting may reduce variations of models and improve the overall performance [64]. Second, although we analyzed the results on three cancer types, data from more cancer types may also be analyzed. Pan-cancer analysis across different cancer types may improve the reliability of our results, and enable us to evaluate the different tumor evolution mechanisms of each cancer type. Third, more advanced methods for survival analysis might be warranted. For example, deep learning has been applied to survival analysis recently [65, 66, 67], demonstrating the feasibility of using a neural network as an effective feature extractor for predicting prognoses if the evolutionary, driver and clinical features of all the available samples are integrated properly.

## 4 Conclusions

Predicting progression outcomes of cancers is a difficult problem underlying many important clinical decisions, such as whether to pursue aggressive treatment, which treatment options to consider, and how frequently to monitor patients for signs of resistance or recurrence. In the present study, we proposed a strategy for enhancing the power for making such predictions through the use of evolutionary features that quantify the degree to which different mutational processes act on a tumor’s genome. We demonstrated via a novel machine learning approach that evolutionary features provide predictive power for future progression and that this predictive power is complementary to that offered by clinical predictors and more conventional “driver-centric” genomic predictors. We further explored the interdependencies among these features and feature classes, showing a complex correlation structure indicative of the heterogeneity in mutational processes across cancers and suggesting how this heterogeneity helps to shape accumulations of driver mutations and ultimately clinical presentations of cancers.

## 5 Methods

### 5.1 Variant calling and evolutionary modeling

For each cancer sample, the somatic genomic variants were first called by a range of possible tools as discussed below, then the SNVs, CNAs and SVs were integrated and converted into a single VCF file, which is the input format required by both evolutionary tree methods considered, Canopy [46] and TUSV [47]. Both methods output fractions of clones in each tumor sample, the inferred phylogenetic tree connecting clones, and the acquired somatic variants of clones during evolution along the tree edges.

We made use of calls from four different variant callers across the three groups of experiments, according to the sequencing strategy of the samples. For the WES BRCA and LUAD samples from TCGA corpus (Table 1 Exp 1), we downloaded SNVs and CNAs from the TCGA Genomic Data Commons Data Portal (GDC). It provided calls from the TCGA pipeline [31], which uses a consensus of standard variant callers such as MuSE [39], MuTect2 [40] and GISTIC2 [41]. For WGS BRCA samples from the TCGA corpus (Exp 2), we called the CNAs and SVs using Weaver [42, 43], while for WGS LUAD samples from TCGA (Exp 2), we called the CNAs and SVs using novoBreak [44]. We did not use phased SVs in both cases for simplicity. Since the number of WGS samples from the TCGA corpus is limited (Exp 2), we made further use of data from the ICGC/PCAWG project [32], which provides a larger corpus of WGS samples for breast cancer and lung cancer (Exp 3). For these samples, we downloaded SNV, CNA, and SV calls^1^, which had been computed for this project using the Sanger pipeline [45]. We pooled the two subtypes of lung cancer: LUAD and LUSC into a single LUCA category to increase the size of the dataset.

We applied two general approaches to derive tumor phylogenetic trees, based on the availability of WGS and WES data. WGS data provides a much better ability to call SVs. To assess the predictive value of SVs, we sought to capitalize on this capability by building phylogenies using a customized version of the TUSV phylogeny software [47], which infers phylogenies from SVs and CNAs and is, to our knowledge, the only tumor phylogeny program currently able to incorporate SVs into its trees. For WES data, we instead used the third-party tool Canopy [46], which makes inferences from SNVs and CNAs.

We selected Canopy because it makes use of SNVs and CNAs and can make inferences from common VCF files. Canopy [46] infers subclones and predicts phylogenies based on an input VCF file specifically combining SNV and CNA data for inference, making it suitable for WES data. First, the clonal decomposition is explored by Markov chain Monte Carlo (MCMC) simulation, and assessed based on the maximum likelihood estimation (MLE). Users need to specify the range for the number of subclones to deconvolve, and also a range for the number of chains and the length of each chain for MCMC simulation. Bayesian information criterion (BIC) is used to determine the optimal number of clones in the specified range. The clonal composition and the bifurcating tree with SNVs and CNAs on the edges are determined based on the posterior distribution. Canopy outputs the clonal compositions and phylogenetic trees, where each SNV or CNA is assigned to a specific edge, which are then extracted as phylogenetic features.

The tumor phylogenetic reconstruction tool TUSV [47] uses a coordinate descent algorithm combining mixed integer linear programming (MILP) to optimize for a minimum evolution CNA model assuming SVs accumulate via a perfect phylogeny. The MILP model implements a trade-off between the likelihood of CNAs described in observed breakpoints and the evolutionary cost of the phyloge-netic tree. We heuristically extended the previously published TUSV for the present work to further incorporate SNVs through a simplified mutation model that excludes the possibility of recurrent mutation but allows for the loss of SNVs through allelic loss. The output of the extended TUSV is a set of inferred clones characterized by the subset of variations inferred in each and a phylogeny connecting those clones.

We finally added a trivial evolutionary model, which we dub the “cumulative evolutionary” model, in contrast to the “phylogenetic evolutionary” model inferred by Canopy or TUSV. This cumulative model is intended as a aggregate approximate model of evolutionary preference derived from overall mutation burdens. For this model, we assume that there is a single branch of evolution from normal to cancer, resulting in a diploid tree of a “normal” root and single derived cancer state. This two-node cumulative evolutionary tree gives a crude approximation to evolutionary rates that requires minimal assumptions about the underlying evolutionary model and data types.

### 5.2 Data processing and feature extraction

Our validation experiments make use of clinical features, assumed to be data that would normally be available for prognosis in clinical practice, and three types of genomic features: driver, cumulative evolutionary features, and phylogenetic evolutionary features. We describe how they were extracted and processed below.

#### Clinical features

All the clinical data for the TCGA samples were extracted from TCGA-reported clinical metadata downloaded from GDC. The clinical features of ICGC/PCAWG samples in Exp 3 also came from GDC, as the samples sequenced using WGS are a subset of the TCGA samples. We removed the features that are available for fewer than half of the samples. We then manually pruned the resulting feature set to provide a consensus representation of information likely to be available to clinicians at the time of diagnosis. The final list of features provided as input to our inference pipeline is shown in Table 7. We note that a large portion of these clinical features is in common between BRCA and LUCA. Examples of preserved clinical features are *person neoplasm cancer status*, *pathologic stage*, and *histological type*.

**Table 7:** List of clinical features. BRCA and LUCA samples share a large portion of similar clinical features. Three data types are available: binary, categorical and continuous.

| Clinical Feature | Data Type | Included |  |
| --- | --- | --- | --- |
|  |  | BRCA | LUCA |
| person neoplasm cancer status | binary | ✓ | ✓ |
| gender | binary | ✓ | ✓ |
| history of neoadjuvant treatment | binary | ✓ | ✓ |
| ethnicity | binary | ✓ | ✓ |
| pathologic stage | categorical | ✓ | ✓ |
| histological type | categorical | ✓ | ✓ |
| race | categorical | ✓ | ✓ |
| age at initial pathologic diagnosis | continuous | ✓ | ✓ |
| lab procedure her2 neu in situ hybrid outcome type | binary | ✓ |  |
| breast carcinoma progesterone receptor status | categorical | ✓ |  |
| breast carcinoma estrogen receptor status | categorical | ✓ |  |
| lab proc her2 neu immunohistochemistry receptor status | categorical | ✓ |  |
| margin status | categorical | ✓ |  |
| her2 immunohistochemistry level result | categorical | ✓ |  |
| number of lymphnodes positive by he | continuous | ✓ |  |
| cytokeratin immunohistochemistry staining method micrometastasis indicator | continuous | ✓ |  |
| menopause status | categorical | ✓ |  |
| her2 neu chromosone 17 signal ratio value | continuous | ✓ |  |
| her2 immunohistochemistry level result | continuous | ✓ |  |
| her2 erbb pos finding cell percent category | continuous | ✓ |  |
| fluorescence in situ hybridization diagnostic procedure chromosome 17 signal result range | continuous | ✓ |  |
| lung cancer type | binary |  | ✓ |
| anatomic neoplasm subdivision | categorical |  | ✓ |

We next preprocessed, encoded, and imputed the clinical features according to their value types. For continuous values, missing values were filled with the median value of the cohort, then were normalized to have zero mean and unit variance. For binary values, missing values were filled with the mode of the cohort, and the features were encoded by 0/1. For non-binary categorical values, missing values were filled with the mode, and a feature of *k* categories was mapped into *k* mutually exclusive binary features.

#### Driver features

The potential drivers of BRCA and LUCA came from two sources. First, we used the IntOGen database [29], where the top 20 drivers based on mutation counts in samples of each cancer type were collected. Second, we used the COSMIC database [56], where 20 common drivers were collected. See Table 8 for the full list of potential drivers of both BRCA and LUCA. We counted the times that a driver was perturbed by SNVs, CNAs or SVs. Examples of common drivers are *TP53*, *PIK3CA*, and *GATA3*.

**Table 8:** List of driver features. The potential drivers come from both IntOGen [29] and COSMIC [56] databases. BRCA and LUAD share a large portion of drivers. Somatic mutation rates of SNV, CNA and SV in all drivers are in continuous value.

| Driver Feature | Included |  |
| --- | --- | --- |
|  | BRCA | LUCA |
| *TP53, PIK3CA, GATA3, MLL3, CDH1, NCOR1, MAP2K4, PTEN, AKT1, RUNX1, NF1, RB1, ARID1A, TBX3, MLL2, SPEN, LRP1B, ESR1, KMT2C, KMT2D, FOXA1, ERBB2* | ✓ | ✓ |
| *MAP3K1, MACF1, MED12, ATM, AKAP9* | ✓ |  |
| *KEAP1, CDKN2A, KRAS, EGFR, STK11, KDR, FAT1, SVEP1, NFE2L2, FN1, NOTCH1, MLL* |  | ✓ |

#### Cumulative evolutionary features

The cumulative features of samples were extracted from overall mutation burdens subdivided by mutation class, analyzed as if they were derived from two-node evolutionary trees (Table 9). Two types of features were included. First, we used mutation rates for different types of common mutations, e.g., *A* → *T*. Because of the relatively sparse data, we did not break these down further into trinucleotide context, as is typically done in mutation signature analyses [12]. Second, we estimated aggregate mutation rates of SNVs, CNAs and SVs. We also took into account the size of CNA region and duplication/deletion of the CNA, e.g., rates of CNA above 500,000 nt (*cnv lg rate*), CNA deletion rates (*cnv del rate*) etc. In each case, we treated mutations as occurring on a single evolutionary tree edge spanning from normal to tumor. When the normal state is unknown, we screened out sites of common germline single nucleotide polymorphisms (SNPs) using germline SNP data from the 1000 Genomes Project [68], and used deviation from a standard human reference (GRCh38/hg38 for TCGA, GRCh37/hg19 for ICGC) [69]. We assumed a fixed edge length of ten years as an estimated time from the appearance of the first ancestral tumor cell to the time of sequencing in order to provide a scaling factor to convert mutation counts into estimated rates, although we note that the scale is arbitrary and does not affect the machine learning inference.

**Table 9:** List of cumulative evolutionary features. The mutation rates related to SNVs, CNAs and SVs of samples are included. All cumulative evolutionary features are in continuous value.

| Cumulative Feature | Definition | Included |  |
| --- | --- | --- | --- |
|  |  | WES | WGS |
| A→T | mutation rates | ✓ | ✓ |
| A→G | mutation rates | ✓ | ✓ |
| A→C | mutation rates | ✓ | ✓ |
| T→A | mutation rates | ✓ | ✓ |
| T→G | mutation rates | ✓ | ✓ |
| T→C | mutation rates | ✓ | ✓ |
| G→A | mutation rates | ✓ | ✓ |
| G→T | mutation rates | ✓ | ✓ |
| G→C | mutation rates | ✓ | ✓ |
| C→A | mutation rates | ✓ | ✓ |
| C→T | mutation rates | ✓ | ✓ |
| C→G | mutation rates | ✓ | ✓ |
| snv_rate | total SNV rates | ✓ | ✓ |
| cna_rate | total CNA rates | ✓ | ✓ |
| cna_amp_rate | CNA duplication rates | ✓ | ✓ |
| cna_del_rate | CNA deletion rates | ✓ | ✓ |
| cna_lg_rate | rates of CNA above 500,000 nt | ✓ | ✓ |
| cna_sm_rate | rates of CNA below 500,000 nt | ✓ | ✓ |
| sv_rate | total SV rates |  | ✓ |

#### Phylogenetic evolutionary features

After a phylogenetic evolutionary tree was built for WES by Canopy, or for WGS by the extended TUSV, measures that quantify topological features of the tree were extracted. As we can see from Table 10, since the outputs of TUSV include extra SV information, we have some additional phylogenetic features for WGS samples. However, there are still some common tree features, such as the clone number (*num clone*), the height of the phylogeny (*height*) and the average of edge lengths (*branch mean*) that are conserved between phylogeny inference methods. All the evolutionary features, cumulative and phylogenetic, were normalized to have zero mean and unit variance before fed into the machine learning module.

**Table 10:** List of phylogenetic evolutionary features. Due to the different output of Canopy [46] (phylogenetic model for WES) and TUSV [47] (phylogenetic model for WGS), the sets of phylogenetic features are slightly different. The WGS data contain additional features related to CNA and SV rates.

| Phylogenetic Feature | Definition | Included |  |
| --- | --- | --- | --- |
|  |  | WES | WGS |
| num_clone | clone number | ✓ | ✓ |
| diversity | diversity of clone portions |  | ✓ |
| lg_clone_portion | portion of the largest clone |  | ✓ |
| lg_clone_snv | SNV rates in the largest clone | ✓ |  |
| lg_clone_cna | CNA rates in the largest clone | ✓ | ✓ |
| lg_clone_sv | SV rates in the largest clone |  | ✓ |
| height_topology | topological height of phylogeny |  | ✓ |
| height | height of phylogeny | ✓ | ✓ |
| height_cna | height of phylogeny in unit of CNA rates |  | ✓ |
| height_sv | height of phylogeny in unit of SV rates |  | ✓ |
| branch_num | number of edges in phylogeny |  | ✓ |
| branch_len | total edge lengths |  | ✓ |
| branch_mean | average of edge lengths | ✓ | ✓ |
| branch_mean_cna | average of edge lengths in unit of CNA rates |  | ✓ |
| branch_mean_sv | average of edge lengths in unit of SV rates |  | ✓ |
| branch_max | maximum edge length | ✓ | ✓ |
| branch_max_cna | maximum edge length in unit of CNA rates |  | ✓ |
| branch_max_sv | maximum edge length in unit of SV rates |  | ✓ |
| branch_var | variance of edge lengths | ✓ | ✓ |
| branch_var_cna | variance of edge lengths in unit of CNA rates |  | ✓ |
| branch_var_sv | variance of edge lengths in unit of SV rates |  | ✓ |

### 5.3 Feature selection

After extracting the four types of features (Sec. 5.2), we implemented a two-stage feature selection method for the machine learning regression model, which consists of a filtering stage and a step-wise forward selection stage.

#### Feature filtering

We first implemented feature filtering to remove features that do not contribute to the prognoses and those that are highly correlated [70, 71]. This can effectively reduce the computational complexity and the hypothesis space to prevent overfitting at the step-wise selection stage [72], and to prevent the multicollinearity across features [73]. Two rules were applied. First, **max-relevance** of the response and individual features: we performed univariate Cox regression analysis [74] on each single feature to find the ones that are related to the prognoses [73]. We did not use more standard simple metrics, such as Bayesian error [70], because the problem of predicting clinical outcomes from censored temporal data, such as survival or recurrence, is very different from a conventional uncensored classification problem. Second, **min-redundancy** of features: we removed one of the features if two features of the same type are highly correlated. In practice, two features with Pearson coefficient in absolute value beyond 0.8 would be taken as highly correlated [73]. Section 2.2 shows that overall the clinical and genomic features are independent of each other and have little correlation, indicating the orthogonal information they entail separately. However, the strong correlations across features within each feature type may lead to multicollinearity and redundancy of the model.

#### Step-wise selection

After the initial feature filtering stage, the top five features plus the ones deemed statistically significant for each feature type were then used for step-wise feature selection. For BRCA in Exp 2, since we have too few samples, we restricted the number of filtered features to be within five. We implemented the forward selection of clinical, driver, cumulative evolutionary, and phylogenetic evolutionary features using 10-fold CV in Exp 1, and LOOCV in Exp 2 and 3, with the optimization goal of maximizing the concordance index (CI) of predictions on the validation sets [75, 76]. Specifically, clinical features were selected step-wise to maximize CI, and then driver features or evolutionary features were selected step-wise to maximize CI. At the evaluation stage, Both CI and logrank test [77] were calculated and conducted for comparison of different sets of features. See Sec. 5.4 for details of regression method, CI and logrank test.

### 5.4 Cox regression and evaluation

The clinical prognostic outcomes of cancer patients, such as OS and DFS, are censored data, meaning that the time to a death event or recurrence event was not observed for some samples due to the limited follow-up time. Therefore, instead of conventional classification or regression methods, we performed Cox regression [74] and evaluated the prediction performance with metrics for survival analysis, which are specifically designed to cope with these censored data. In our formulation, the samples are

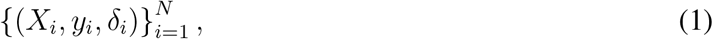

where *N* is the total number of samples, *X*_*i*_ ∈ ℝ^*m*^ is the feature vector of sample *i*, *δ*_*i*_ ∈ {0, 1} indicates the status of patient *i* at the last follow-up time *y*_*i*_ = min (*T*_*i*_, *C*_*i*_): If *δ*_*i*_ = 1, the event happened and was observed at time *y*_*i*_ = *T*_*i*_. If *δ*_*i*_ = 0, the event had not happened at the censoring time *y*_*i*_ = *C*_*i*_.

Cox regression is a semi-parametric regression method based on the proportional hazards (PH) assumption:

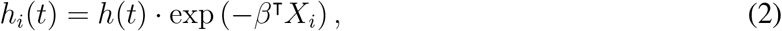

where *h*_*i*_(*t*) ≔ lim_Δ*t*→0_ Pr(*t* < *T*_*i*_ ≤ *t* + Δ*t* | *T*_*i*_ > *t*)/Δ*t* is the hazard of patient *i* at time *t*, or in another word, the probability of death if the patient has survived to time point *t* (for OS; similarly it is the hazard of recurrence for DFS), *h*(*t*) is the non-parametric part calculated from the training data, *X*_*i*_ is the feature vector of sample *i*, *β* ∈ ℝ^*m*^ is the model parameter to be estimated. Instead of predicting whether the patient will be dead or alive, Cox regression estimates *β* and thus provides the hazard of the patient following Eq. (2). When the total number of samples is large, we can roughly assume *h*(*t*) to be close enough at the time of CV. The comparison of *h*_*i*_(*t*) thus reduces to the comparison of risk score *η*_*i*_ = − *β*^T^*X*_*i*_, i.e., the logarithm of “hazard ratio” exp (−*β*^T^*X*_*i*_), which is independent of time *t* [78].

At the time of training, we optimized the negative log-partial likelihood function of Cox model:

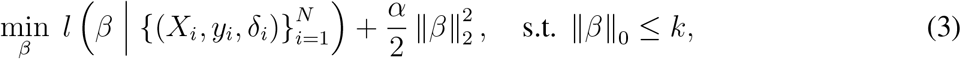

where *α* is chosen to be a small number here mainly for the stability during optimization and prevent overfitting, || *β* ||_0_ is the *ℓ*_0_-norm that counts the number of nonzero coefficients in *β*. The parameter *k* is chosen to maximize the prediction performance on the validation set.

With the predictions of risk scores, we can evaluate prediction results based on an assessment of concordance index (CI) [75, 76], and an assessment of the statistical significance of separating censored survival data using a logrank test [77]. The CI is defined as the following:

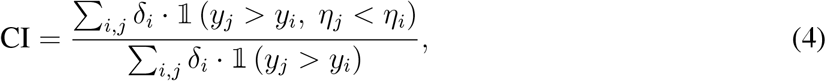

where 𝟙 (statement) is the indicator function. CI is a value similar to area under curve (AUC). A model reaches perfect prediction when CI = 1 and random guess when CI = 0.5. The logrank test uses a statistical test to accept or reject the null hypothesis 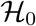: two groups of samples share the same survival profile. It calculates a statistic *z*^2^ from observations of two groups of censored data. While 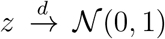, we can get the *p*-value to accept or reject 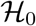. In our experiments, we classified all samples into two groups with the median of risk scores 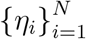, and used logrank test to evaluate the differences between these two cohorts of predicted malignant and benign samples. The Cox regression and logrank test were implemented using Python package `lifelines` [79].

## Supporting information

Supplemental Figures of Feature Correlation

## Declarations

### Availability of data and materials

The source code for the pipeline during the current study is available in the Github repository.^2^ The public data of TCGA and ICGC analyzed during the current study are freely available for download from the respective consortia. The controlled data that support the findings of this study, such as raw sequencing data are available from TCGA and ICGC although restrictions apply to the availability of these data, which were used under license for the current study, and so are not publicly available.

### Authors’ contributions

R.S. conceived general ideas, supervised implementation, planned validation, and interpreted experimental results. J.M. conceived general ideas, supervised aspects of implementation and validation and the interpretation of experimental results. Y.T. developed new machine learning methods for analysis pipeline, and implemented and executed validation experiments. A.R. developed and implemented aspects of method validation, and advised others on the use of variant calling. X.C. integrated phylogenetic methods into the pipeline, and contributed to the design and execution of validations studies. Z.C. implemented feature extraction and selection methods for analysis pipeline, and contributed to the design and execution of validation experiments. J.E. and H.K. developed and implemented phylogenetic methods for the pipeline, and contributed to the design and execution of validation experiments. All authors contributed to writing, reviewing, and/or editing the manuscript.

## Acknowledgements

The results published here are in whole or part based upon data generated by TCGA managed by the NCI and NHGRI. Information about TCGA can be found at http://cancergenome.nih.gov. We likewise thank the ICGC for additional data on which these results were in part based. We would like to thank Bjarni V. Halldörsson for his insightful comments and suggestions and Cenk Sahinalp and Ben Raphael for helpful guidance.

## Footnotes

1 https://dcc.icgc.org/repositories. Accessed 6 February 2019.

2 https://github.com/CMUSchwartzLab/cancer-phylogenetics-prognostic-prediction

