## Supplemental Figures of Feature Correlation for "Improving personalized prediction of cancer prognoses with clonal evolution models"

### Supplemental Information

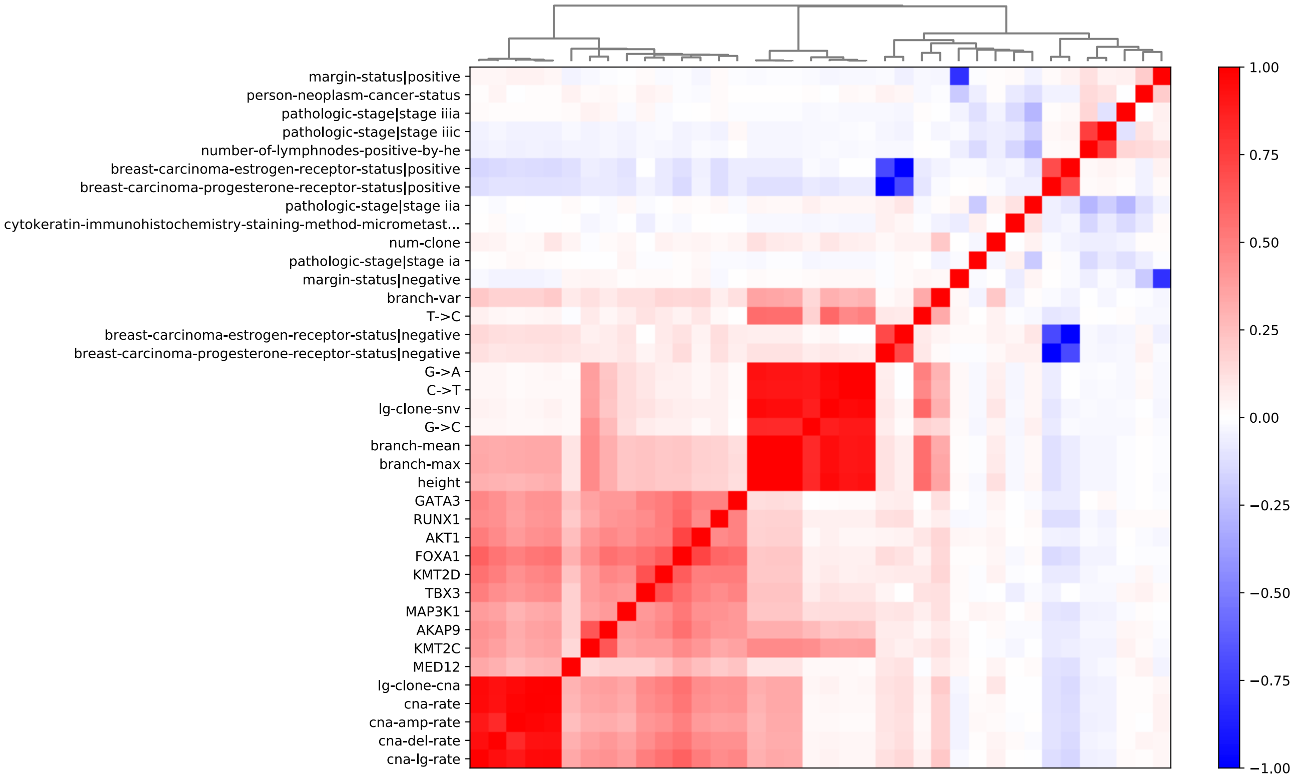


**Figure S1: Correlation matrix of features (Exp 1 BRCA DFS).**

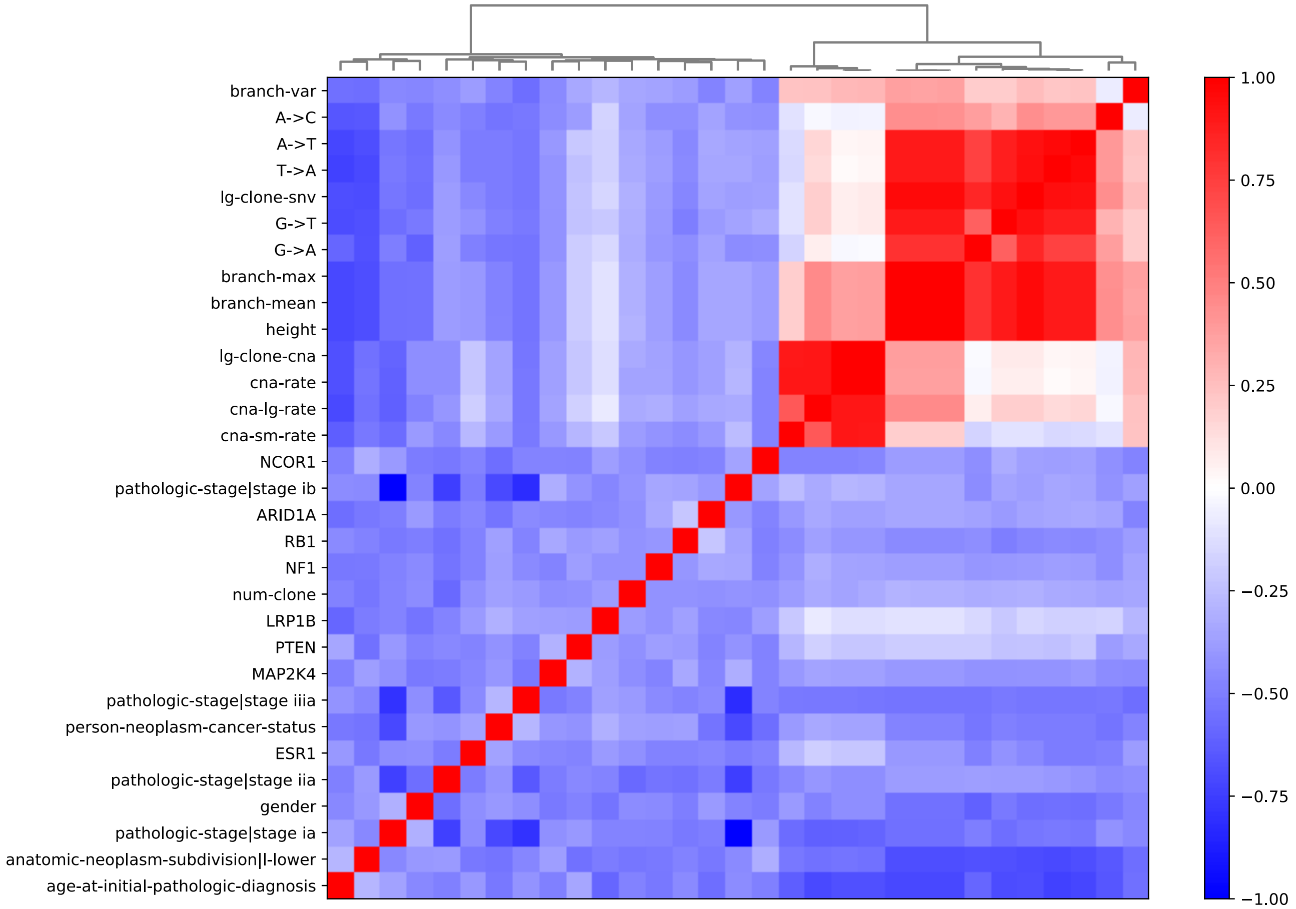


**Figure S2: Correlation matrix of features (Exp 1 LUAD OS).**

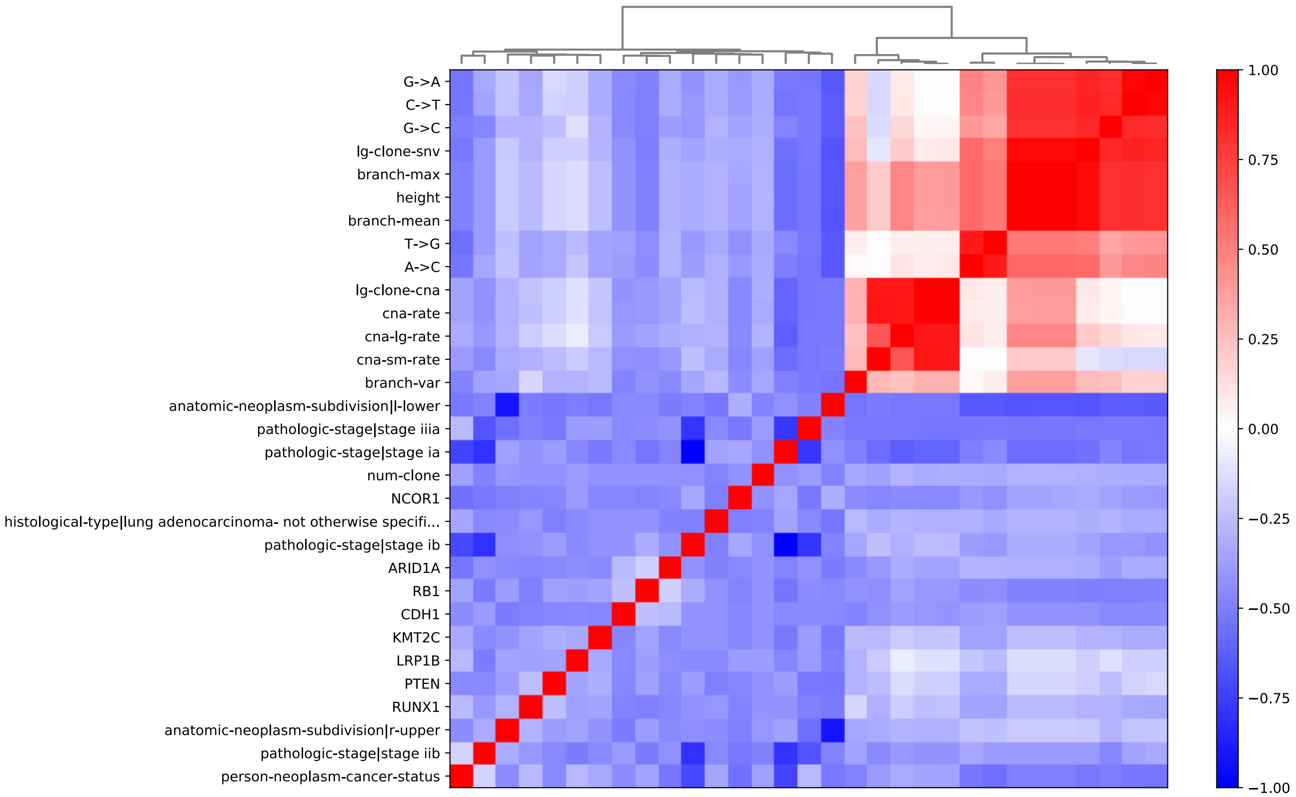


**Figure S3: Correlation matrix of features (Exp 1 LUAD DFS).**

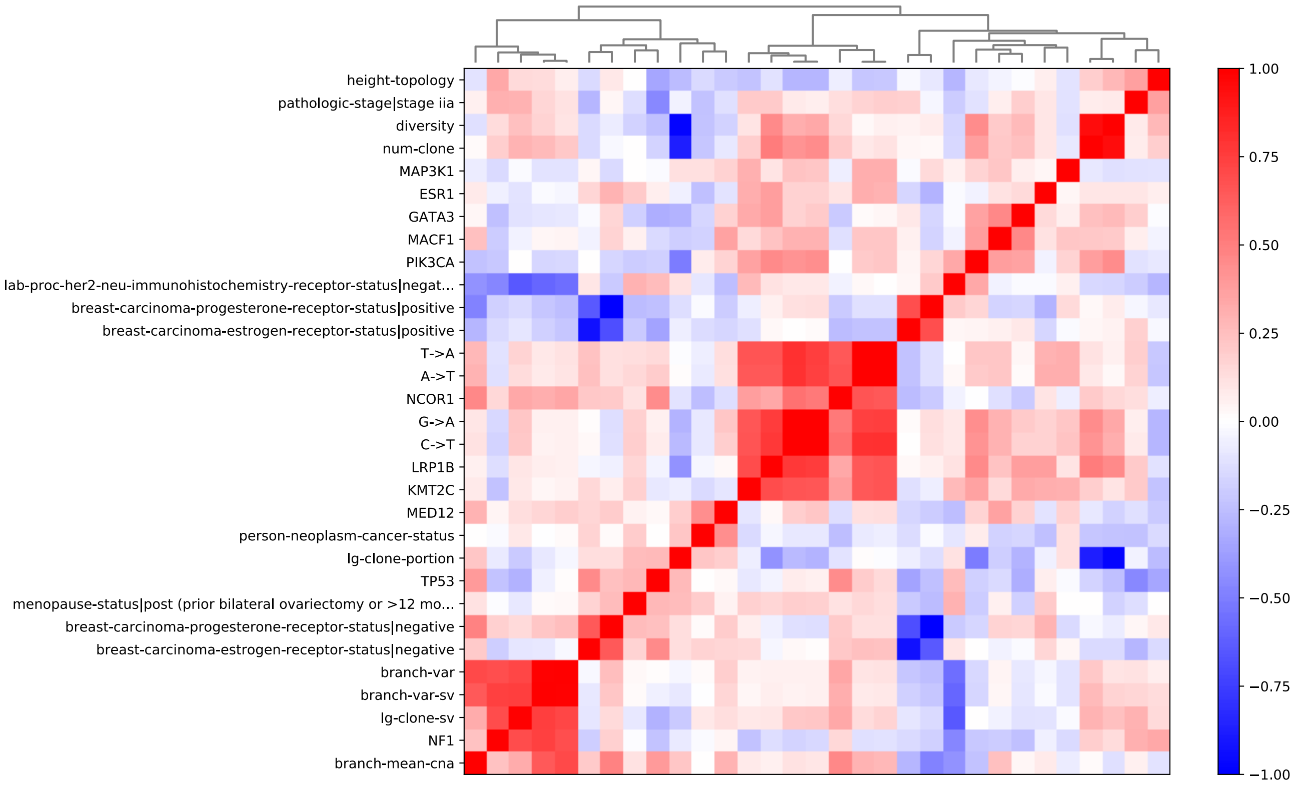


**Figure S4: Correlation matrix of features (Exp 2 BRCA OS).**

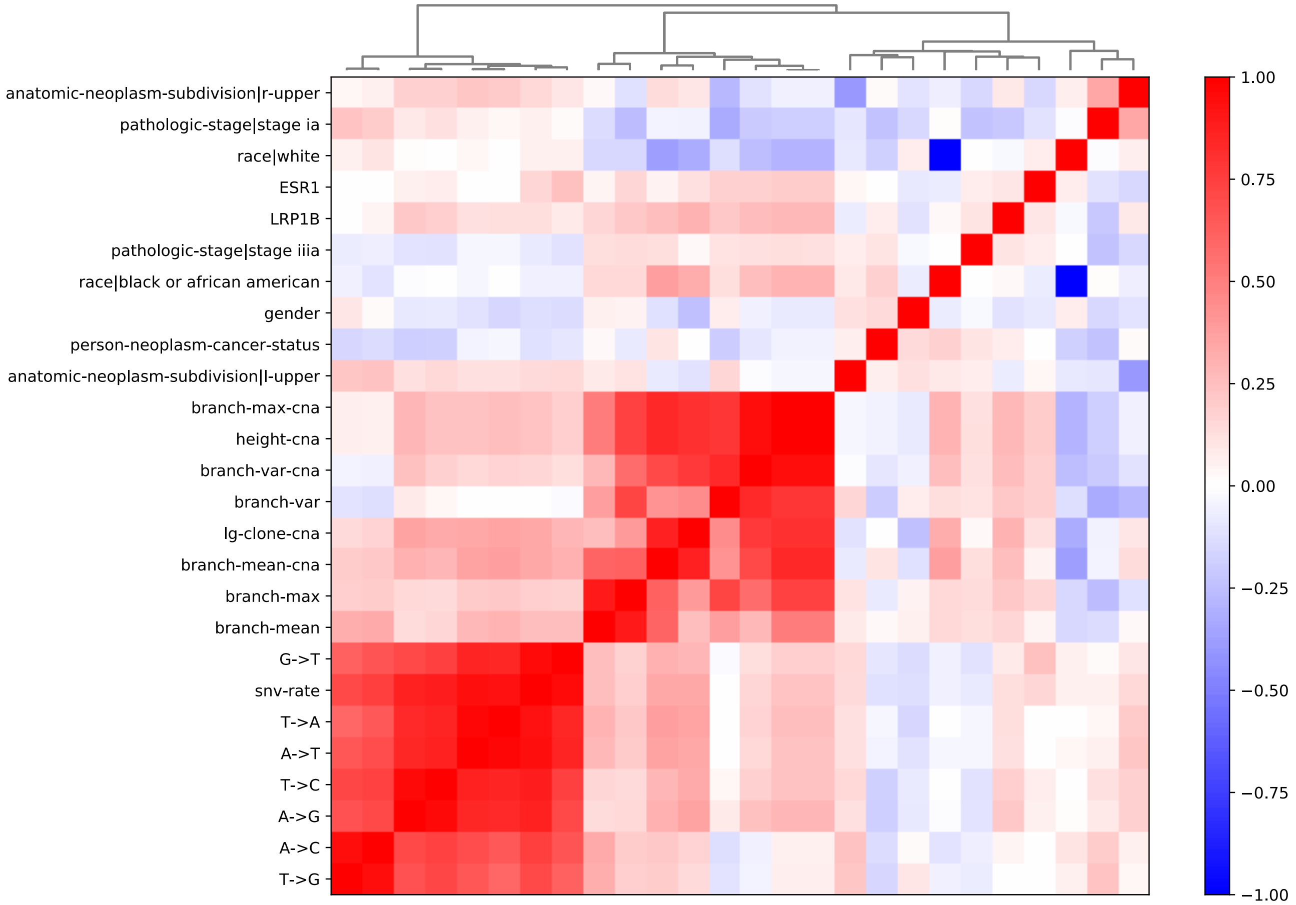


**Figure S5: Correlation matrix of features (Exp 2 LUAD OS).**

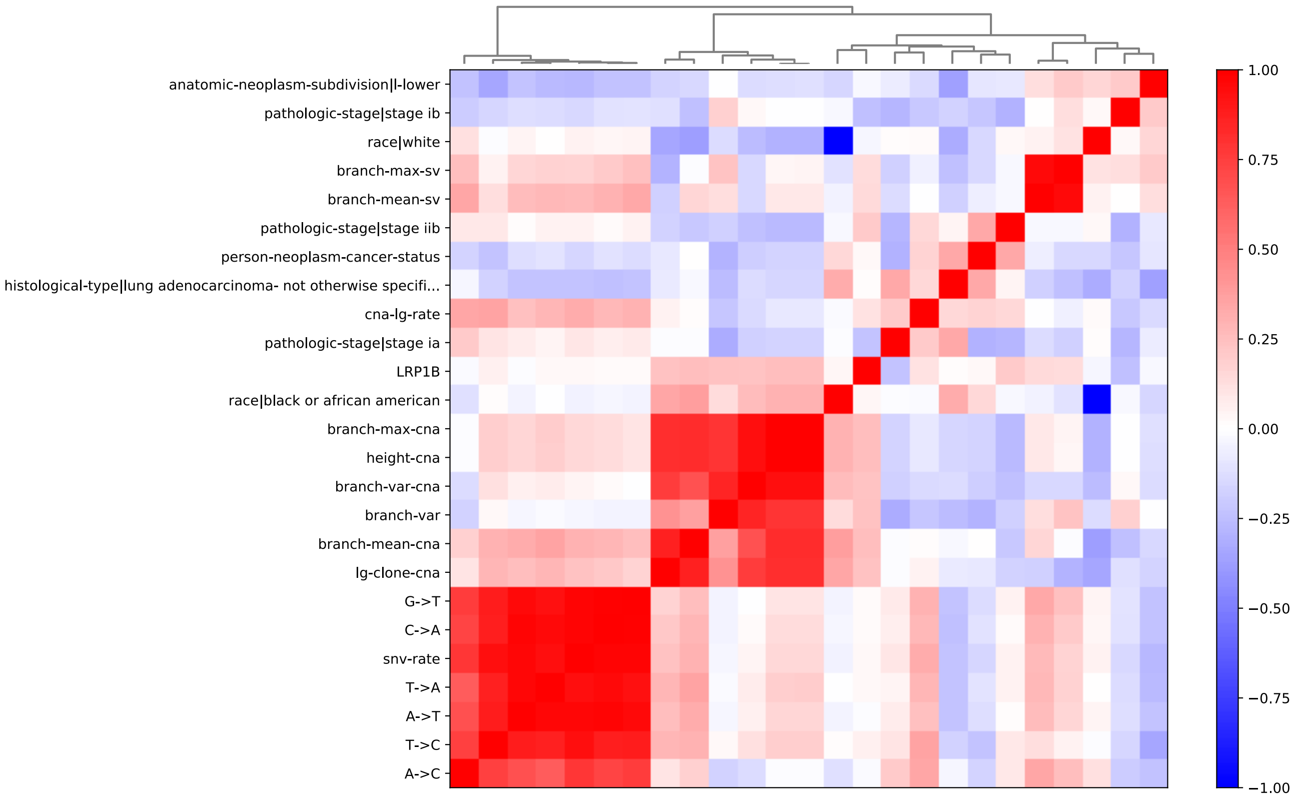


**Figure S6: Correlation matrix of features (Exp 2 LUAD DFS).**

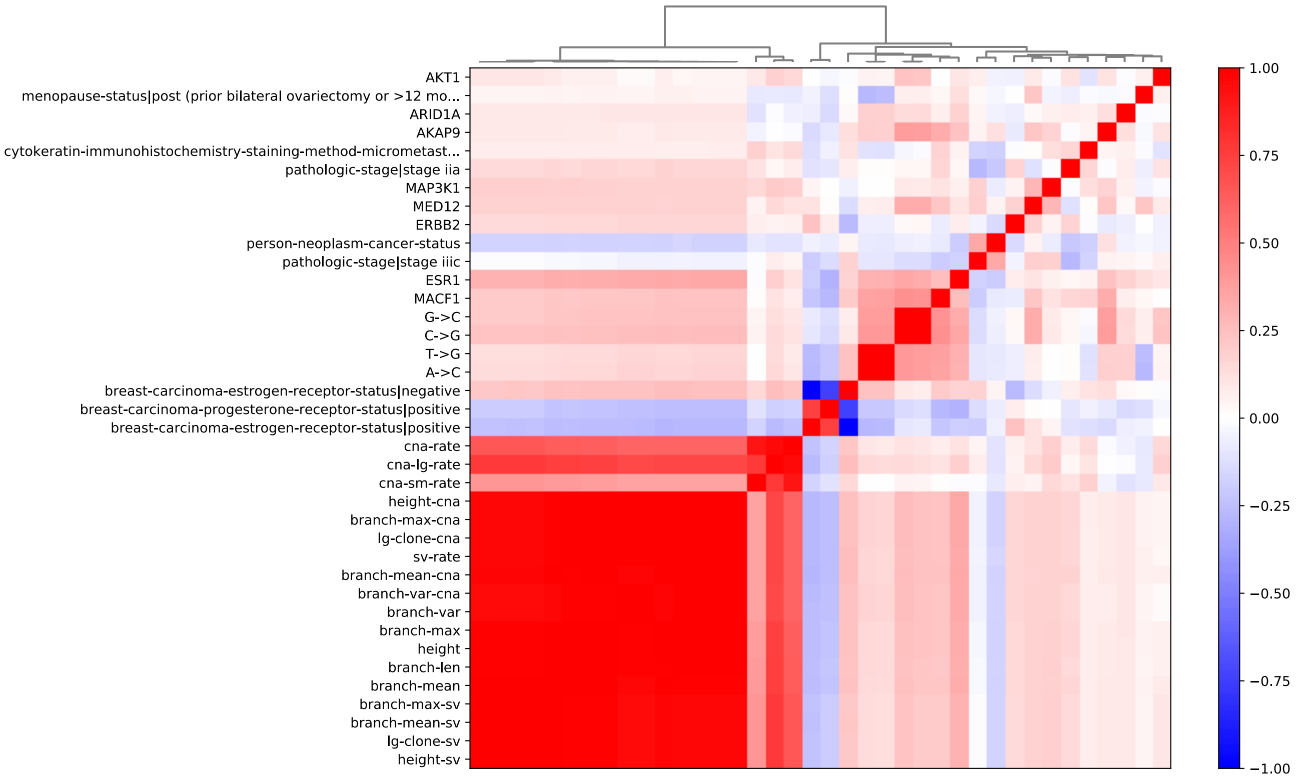


**Figure S7: Correlation matrix of features (Exp 3 BRCA OS).**

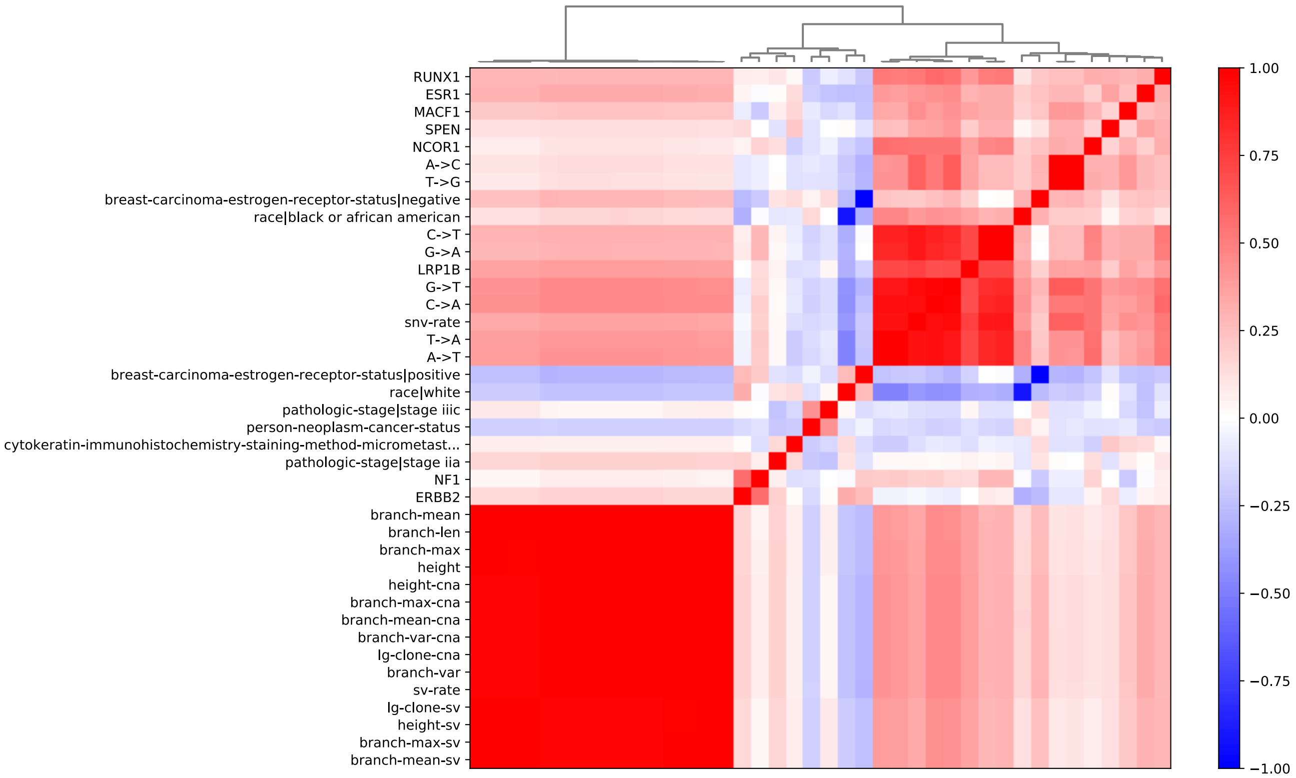


**Figure S8: Correlation matrix of features (Exp 3 BRCA DFS).**

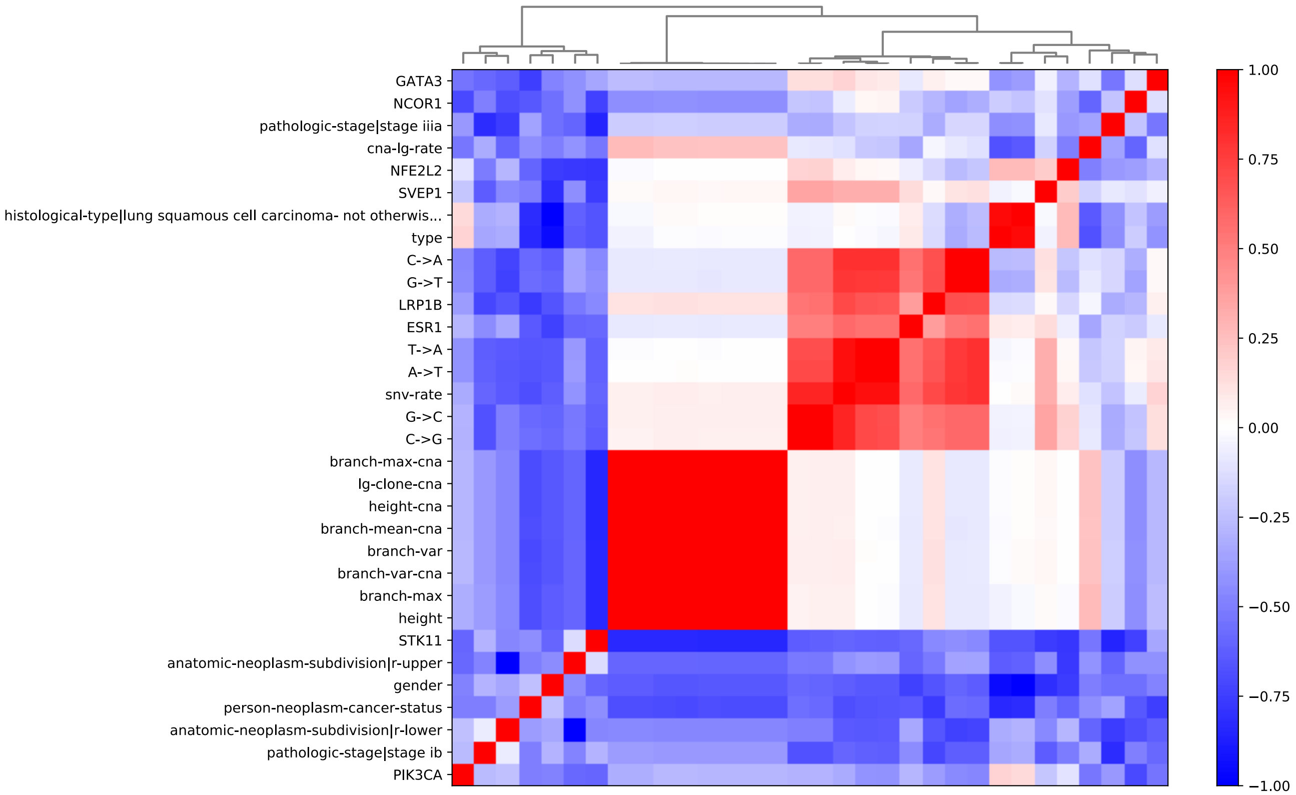


**Figure S9: Correlation matrix of features (Exp 3 LUCA OS).**

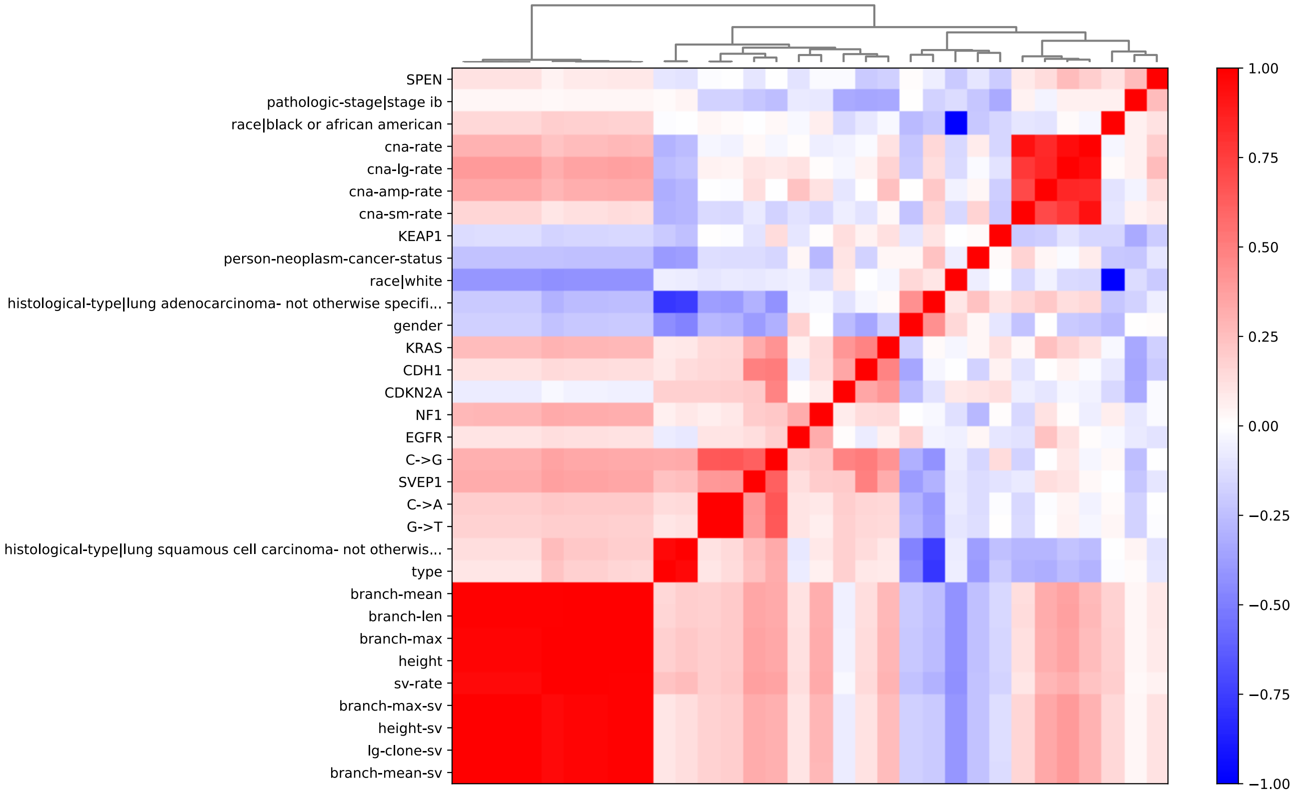


**Figure S10: Correlation matrix of features (Exp 3 LUCA DFS).**
